# Breaking Down Cell-Free DNA Fragmentation: A Markov Model Approach

**DOI:** 10.1101/2023.07.06.547953

**Authors:** Terence H.L. Tsui, Phil F. Xie, Salvador Chulián, Víctor M. Pérez-García

## Abstract

Cell-free DNA (cfDNA) is released into the bloodstream from cells in various physiological and pathological conditions. The use of cfDNA as a non-invasive biomarker has attracted attention, but the dynamics of cfDNA fragmentation in the bloodstream are not well understood and have not been previously studied computationally.

To address this issue, we present a Markovian model called FRIME (Fragmentation, Immigration, and Exit) that captures three leading mechanisms governing cfDNA fragmentation in the bloodstream. The FRIME model enables the simulation of cfDNA fragment profiles by sampling from the stationary distribution of FRIME processes. By varying the parameters of our model, we generate fragment profiles similar to those observed in liquid biopsies and provide insight into the underlying biological mechanisms driving the fragmentation dynamics.

To validate our model, we compare simulated FRIME profiles with mitochondrial and genomic cfDNA fragment profiles. Our simulation results are consistent with experimental and clinical observations and highlight potential physicochemical differences between mitochondrial and genomic cfDNA.

The FRIME simulation framework provides an initial step towards an improved computational understanding of DNA fragmentation dynamics in the bloodstream and may aid in the analysis of liquid biopsy data.

**Author summary:** Cell-free DNA (cfDNA) released into the bloodstream from cells in different conditions is the basis for liquid biopsies, thus being non-invasive biomarkers of potential interest. However, the dynamics of cfDNA fragmentation in the bloodstream remain poorly understood and are yet to be studied computationally. To address this issue, we developed a Markovian model which captures the three leading mechanisms governing cfDNA fragmentation in the bloodstream: FRagmentation, IMmigration, and Exit (FRIME). The FRIME model enables the simulation of cfDNA fragment profiles by sampling from the stationary distribution of FRIME processes. Simulation results are compared with mitochondrial and genomic cfDNA fragment profiles, which show consistency with experimental and clinical observations. The FRIME simulation framework is a significant step towards understanding DNA fragmentation dynamics in the bloodstream, uncovering potential physicochemical differences between mitochondrial and genomic cfDNA, and aiding the analysis of liquid biopsy data.

## Introduction

Cell-free DNA (cfDNA) is extracellular DNA present in biofluids, particularly blood, that originates mainly from dying cells as part of normal cellular turnover or pathological processes, including cancer [1]. Assays, such as PCR and sequencing, can detect cfDNA in the bloodstream, providing clinicians with molecular information on the cell-of-origin. This created a new field called “liquid biopsy”, which is more accessible and less risky than traditional biopsies, which require invasive procedures that may cause complications such as infection, bleeding, or pain. The decreasing cost of sequencing technologies has made cfDNA a promising material for prenatal genetic diagnosis, early cancer detection, and cancer treatment decision-making [2]. Currently, over 1000 clinical trials are exploring the potential of cfDNA, and there are a few FDA-approved tests [3].

Apart from being low-risk and easier to obtain, cfDNA can provide real-time monitoring of the disease status, allowing for prompt therapy adjustments. However, several challenges must be addressed before cfDNA can become a standard clinical tool. These include the need for further validation of the methods used for cfDNA detection, identification of the most appropriate clinical applications, optimization of the cost-effectiveness of cfDNA analysis and an improved understanding of the processes behind cfDNA generation and dynamics [4].

There are many molecular features in cfDNA data, such as DNA mutation, copy number variation [5], DNA methylation [6], and histone modifications [7]. When focusing on the oncological applications of cfDNA, identifying pathological signals from physiological activities or technical background in cfDNA is non-trivial. This is because tumor-derived cfDNA usually makes up less than 10% of plasma cfDNA even at late tumor stages [8], resulting in low signal-to-noise ratio. Sometimes, tumor cfDNA cannot be detected despite using highly sensitive methods such as digital PCR [9]. Strategies to overcome this problem include enriching for tumor-derived DNA [10], as well as combining multi-modal information to increase the number of tumor-specific features that can be captured or analyzed [11, 12].

CfDNA fragment length profile is one of the keys in both strategies. It has been demonstrated in mice xenograft models and cancer patients that tumor-derived cfDNA are in general shorter and more fragmented [10, 13], therefore selecting for short DNA fragments between 90-150 base pairs (bp) can enrich tumor signals. The short-to-long fragment ratio has also been proposed to be a diagnostic marker [14]. Yet, studies so far have been focusing on fragment sizes under approximately 200bp, and fragment length profiles beyond this have not been thoroughly described. Moreover, how to best integrate the fragmentation profile with other multi-modal information remains an open question.

Currently, cfDNA data is commonly perceived by biologists as a fixed snapshot, with testing for specific markers of interest, such as mutation frequency. Nevertheless, cfDNA is part of a dynamic equilibrium similar to other biological processes, constantly being generated, broken down, and eliminated from the body. Although the kinetics of cfDNA can potentially be analyzed during tumor resection [15] or caesarean section [16], these approaches are not practical for diagnostic or molecular characterization purposes.

Fragmentation patterns have been proposed to have a high potential of containing information of clinical relevance in what it is currently being referred to as cfDNA framentomics. Further insights into the fragmentomics of plasma cfDNA may shed light on the origin and fragmentation mechanisms of cfDNA during pathological processes in diseases and enhance our ability to take the advantage of plasma cfDNA as a molecular diagnostic tool. This includes extracting information from fragment length, end motifs, jagged ends, preferred end coordinates, as well as nucleosome footprints, open chromatin region, and gene expression inferred by the cfDNA fragmentation pattern across the genome [20–22].

However, to unveil the cfDNA properties, including that of fragmentation patterns, as a potential biomarker for early diagnosis, diagnosis and prognosis, it is crucial to characterize the intrinsic kinetics of cfDNA within the organism. This is so especially when using cfDNA analysis as longitudinal diagnostic tool for monitoring the course of treatment [17]. Consequently, computational models that capture the dynamics of cfDNA may serve as a valuable tool to extract biological insights from the static cfDNA data.

Mathematicians and physical chemists have used fragmentation equations to model the degradation of polymers [18, 19]. Fragmentation equations are differential equations that describe how the concentration of fragments at different length scales changes over time. However, these equations did not account for the biological noise that occurs naturally or the advent of DNA sequencing technology. While fragmentation with immigration processes have been studied [23, 24], no mathematical or computational model has been built including an exit mechanism to study the effect of cfDNA clearance mechanisms on the fragment profile. Simple approaches have estimated basic averaged kinetic parameters for different processes [17] but without incorporating the complexities of fragmentation processes. Recent studies based on non-dynamical approaches have correlated tumor detection size with circulating tumor DNA shedding [27] and developed a mathematical equation relating the distribution profile of a stochastically fragmented DNA sample to the probability that a DNA fragment within that sample can be amplified by any PCR assay of arbitrary length [26]. Finally, data analytic mathematical and machine learning method have been used extensively to analyze cfDNA fragmentation datasets [25, 28, 29]. Nevertheless, those approaches have not incorporated mechanistically the dynamical interplay of the different processes playing a role in the fragmentation dynamics.

CfDNA samples collected before and after treatment can provide an expanded view of the genetic response of a patient’s tumor, including the dynamic changes in the mutational landscape as well as the heterogeneity that develops due to the selective pressure of therapy. In addition, it is clear that the potential of cfDNA analyses is enhanced when combined with results obtained from tissue biopsies and integrated with mathematical modelling of tumor evolution. [30].

Therefore, simulation-based computational frameworks are required to gain a deeper understanding of the biological processes that occur during DNA fragmentation and to provide a more robust clinical analysis, an thus develop biomarkers, of sequenced cfDNA fragment profiles.

In this study, we develop a mathematical/computational stochastic Markovian model and propose that the dynamic behaviour of cfDNA can be retrieved from the fragment length distribution of cfDNA, obtained from sequencing. When a long DNA molecule is cleaved by DNase, the resulting shorter molecules remain in the system, which would be further split into even smaller molecules. When the system reaches dynamic equilibrium, the abundance of molecules of each size would be dependent on the fragmentation behaviour.

The structure of this paper is as follows: In the Methods section, we describe the biological factors that determine the dynamics of cfDNA and propose the FRagmentation with IMmigration and Exit (FRIME) framework, along with an algorithm to implement it. In the Results section, we use the FRIME framework to simulate the distribution of cfDNA fragment lengths and test various biological assumptions by adjusting the parameters of the model. We then compare our simulated results with real-world cfDNA data obtained using state-of-the-art sequencing techniques. In the Discussion section, we explore the biological implications of our results. Our simulations elucidate the shape of the entire cfDNA fragmentation profile, including the fragment counts of high molecular weight fragments. Moreover, our study identifies a difference in fragmentation kinetics between genomic and mitochondrial cfDNA, revealing new insights into how cfDNA is metabolized. We also provide a novel framework to study the dynamic parameters of cfDNA from existing data, which may improve analytic strategies. By shedding light on the underlying biological mechanisms of cfDNA fragmentation, our study contributes to the development of cfDNA as a noninvasive biomarker in clinical settings.

## Materials and methods

### Biological determinants of cfDNA dynamics

Various pathways of cell death, degradation, and regulated extrusion, partial or complete genomes of various origins (e.g., host cells, fetal cells, and infiltrating viruses and microbes) contribute to the shedding into human body fluids, most notably blood, of segmented cell-free DNA (cfDNA) molecules [32]. Active and passive release and elimination mechanisms contribute to the composition of cfDNA. In this manuscript we assumed these macromolecules to be continuously produced, metabolized, and removed from blood through the interplay of the following processes:

#### Production

CfDNA originates mainly from dying cells, which can be part of normal cellular turnover or pathological processes such as cancer [1]. When cells die, mitochondrial and genomic DNAs are released into the bloodstream. There are variations in blood cfDNA levels among patients with different tumor types and stages. In general patients with benign lesions or with early-stage cancer have lower amounts of cfDNA compared to patients with advanced or metastatic tumors of comparable size, what could be a result of the increased cell turnover and metabolic properties of progressing cancer [35]. It has been found that cfDNA levels were correlated with metabolic disease volume, estimated with ^18^F-labelled fluorodeoxyglucose positron emission tomography, in some cancers. In our modelling framework, cfDNA input mechanisms, whatever their origin, were incorporated as an input term feeding large fragments sizes into the system.

#### Fragmentation rate

CfDNA can be digested both intracellularly during cell death and extracellularly through different processes. The fragmentation rate of cfDNA depends on the DNase activity, which can differ across different body fluids [31], health conditions [33], and individuals. In the context of plasma cfDNA, it is digested extracellularly by circulating DNases that break partially digested cfDNA into even smaller fragments.

#### Fragmentation pattern

Healthy and diseased cells exhibit a structured genomic DNA arrangement, with nucleosomes tightly binding most of the DNA except for short linker stretches [34]. DNases selectively digest the linker DNA, creating a periodic pattern of fragment sizes, where each peak corresponds to DNA associated with nucleosomes. This pattern is referred to as a nucleosomal pattern. In contrast, mitochondrial DNA is not bound by nucleosomes and lacks a nucleosomal fragmentation pattern.

#### Elimination

The elimination mechanisms of circulating cfDNA are not fully understood. Nucleosome-bound cfDNA is thought to mainly be cleared by the liver, whereas naked single-stranded DNA may be cleared through the kidney in a size-dependent manner [35]. Since the concentration of cfDNA is more or less in a dynamic equilibrium, it can also be inferred that elimination of cfDNA is concentration-dependent. The reported half-life of cfDNA is between 0.25 to 2 hours [15, 16].

### Technical determinants of cfDNA fragmentation data

There are multiple factors that may cause a systematic bias in cfDNA fragment lengths readings, including the choice of cfDNA extraction kit [36], single-stranded versus double-stranded library preparation [37], and most notably, short-read versus long-read sequencing platform. Short-read sequencing platforms are designed for short DNA molecules, and are not ideal for our study for a few reasons. Firstly, the sequencing length is not guaranteed to cover the entire DNA molecule, and fragment length is inferred from the reference genome after sequence alignment. Thus, fragment length inference is affected by alignment errors as well as host-specific genetic variations. Secondly, there is a fragment length bias in short-read sequencing, where large DNA fragments are captured very inefficiently. This may be an effect of PCR during library preparation or the sequencer itself [38]. On the other hand, long-read sequencing is capable of directly measuring the fragment length by reading through the whole DNA molecule. It can also capture large DNA fragments efficiently with a modest size bias [39].

### Computational model

To model the dynamics of cell-free DNA fragments in the bloodstream, we proposed the FRIME framework which integrated the mechanisms mentioned above into a continuous time Markov process.

See Fig 1 for a schematic diagram of the process.

**Fig 1.**
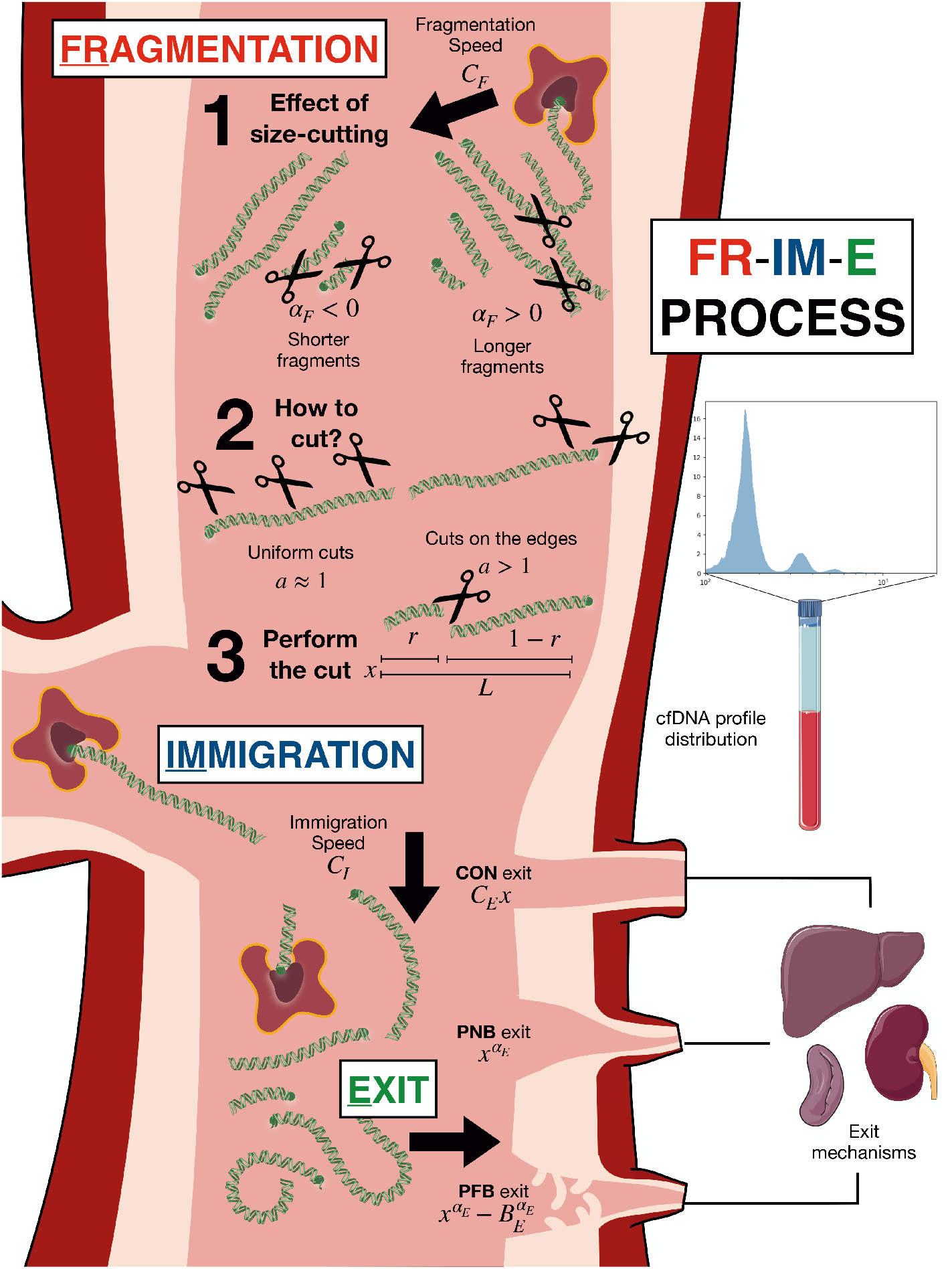
Schematic diagram of the FRIME process of the distribution of cfDNA fragments with length *x*. The FRagmentation mechanisms included the effect of: 1) fragmentation speed *C_F_* and fragmentation power *α_F_* (negativity and positivity related to shorter and longer fragment lengths, respectively); 2) the cut location, with *a* = 1 as uniform cuts, and *a* ≫ 1 as cuts near the edges; 3) the fragmentation ratio *r*, which followed a distribution *Beta*(*a,* 1) for any possible fragment of length *L*. The IMmigration processes were modelled by an immigration speed *C_I_*. For the Exit mechanisms we considered three different possibilities. First a constant exit rate (CON, with *C_E_* as the constant), next a Power decay with No Boundary (PNB, with *α_E_* as the exit rate), and finally a Power decay exit with Fixed Boundary (PFB, with *α_E_* as the exit rate and *B_E_* as the boundary). Icons used are originally from https://bioicons.com, with license https://creativecommons.org/licenses/by/3.0/.

#### FRagmentation mechanisms

CfDNA fragmentation is known to occur through DNase digestion, however it remains unclear how exactly these biomolecules are broken down and how does the reaction rate of cfDNA digestion depend on their size. Reaction rates of other polymers are known to be proportional to polymer sizes raised to some power [40]. Similar results can also be deduced mathematically under the Zimm chain model which considers random movement of polymers [41]. In this paper we considered different types of fragmentation rates and tested their outcomes against human data to see which one provided a faithful description of the observed phenomenology. Such comparison with clinical data can help us find possible biological mechanisms for cfDNA digestion.

We also assumed that a fragment of length *x* reacts with an enzyme at a rate *C_F_ x^α_F_^*. Upon reaction, the fragment breaks into two pieces of length *xr, x*(1 *− r*), where *r ∼ Beta*(*a,* 1). In this paper, the parameters *C_F_, α_F_, r* will denote the *fragmentation speed*, *fragmentation power* and *fragmentation ratio*, respectively.

The case of *α_F_ <* 0 corresponds to an increase of fragmentation rate with shorter fragment lengths. This corresponds to the hypothesis that the reaction rate of cfDNA polymers are limited by diffusion, and longer cfDNA polymers diffuse slower than shorter polymers due to its size and possible secondary structures.

The case *α_F_ >* 0 conversely assumes that fragmentation rate increases with longer fragment length. This scenario corresponds to cfDNA digestion by surface exposure, where the rate would be proportional to the number of available reaction sites.

The parameter *a* determines the distribution of the fragmentation ratio *r*. When *a* = 1, the fragmentation ratio *r* is given by Beta(1,1), and is thus uniformly distributed in [0, 1]. Biologically, this would correspond to the situation where cfDNA is digested by endonucleases, i.e. fragments are cut in the middle. When *a* ≫ 1, the fragmentation ratio *r* is much biased to values close to 1. This last situation would be where the dominant fragmentation mechanism is digestion by exonucleases, i.e. fragments are cut near the edges. It is known that the major intracellular and extracellular DNases responsible for cutting cfDNA are not exonucleases [1, 42], thus in we chose *a* = 1. As described in the paragraph *Fragmentation pattern*, genomic DNA is nucleosome-protected, and fragmentation is not uniform. However, we still assumed uniform cutting on a nucleosomal scale.

#### IMmigration mechanisms

We assumed that the maximum possible length of an immigrated fragment is *L*, that would be equal to or smaller than the size of a full DNA molecule. Fragments of all sizes between [0, *L*] immigrate into the system with rate *C_I_*. As there was no prior knowledge on the length distribution of freshly released cell-free DNA, the length of new fragments were assumed to be uniformly distributed in the interval [0, *L*]. The parameter *C_I_* was chosen to denote the *Immigration speed*.

An alternative is to assume uniform cutting by intracellular DNases of DNA before entering the bloodstream. This scenario would result in more short fragments than long fragments, and could be modelled using a *Beta*(1, *n*) distribution for the immigrating fragment size distribution. The stationary fragment distribution profiles under such an assumption were also studied in this manuscript and described in detail in Section 1 of the supplementary material S1 Appendix.

#### Exit mechanisms

We considered three different classes of exit mechanisms: a constant exit rate (CON), Power decay with No Boundary (PNB), and a Power decay exit model with a Fixed Boundary (PFB).

For the CON model, we assumed that fragments of all sizes exit at the same rate *C_E_*. This implies that cfDNA clearance mechanism is agnostic to cfDNA size. The PNB model corresponds to the assumption that fragments of length *x* exit the system at a rate proportional to *x^α_E_^* with *α_E_ <* 0. That functional dependence describes situations where all fragments leave the system but smaller fragments are easier to release from circulation. Finally, the PFB model is based on the assumption that only fragments smaller than *B_E_* exit the system and do so at a rate proportional to 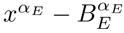. This modelling scenario is in-line with the the renal clearance hypothesis, where only short cell-free DNA fragments can pass through the glomerular capsules and be cleared.

In the following sections of this paper, the parameter *α_E_* is referred as the *exit power* and the parameter *B_E_* as the *exit boundary*. A summary of all three exit mechanisms to be studied in this paper can be found in Table 1. One of our goals in this paper was to study which mechanism is more likely to be the one responsible for cfDNA clearance.

**Table 1.** Functional dependencies of the exit rates *ε* (*x*) on the fragment sizes *x* for the different exit mechanisms considered in this paper.

| Exit Mechanism | $\mathcal{E}(x)$ |
| --- | --- |
| CON | $C_E$ |
| PNB | $L^{-\alpha_E} x^{\alpha_E}, \alpha_E < 0$ |
| PFB | $L^{-\alpha_E} \max\{x^{\alpha_E} - B_E^{\alpha_E}, 0\}$ |

#### FRIME Process and equilibrium measure

To model cfDNA fragmentation profile, a Markov model with the above fragmentation, immigration, and exit mechanisms was studied. Fragment profiles were sampled from the equilibrium measure of the model. More specifically, Monte Carlo simulations were run for a continuous-time Markov Chain (FRIME process) satisfying the following properties:

- The process starts off with a fragment with length *x_i_*(0) ∼ *U* (0, *L*).
- Fragments of length *x* at rate *C_F_ x^α_F_^* fragment into two pieces of length *xr* and *x*(1 *− r*), where *r ∼ Beta*(*a, b*).
- Fragments of length *x < B_E_* leave the system at rate *ε* (*x*).
- New fragments immigrate into the system as a Poisson process with intensity *C_I_*. The length *z* of a new fragment is given by the uniform distribution between [0, *L*].

### Algorithm

The following algorithm was used to generate one simulation of a stationary profile of the FRIME process.

1. Generate *x*_0_ ∼ *U* (0, *L*). Assign ℒ = [*x*_0_]. Assign *ℒ^hist^* = ℒ. Assign *d* = 1, *T* = 0.
2. While *d < κ*:

a. Generate *t_imm_* ∼ *Exp*(*C_I_L*), *t_frag_* ∼ *Exp*(∑_*x*∊ℒ_ *C_F_x^α_F_^*) and *t_E_ ∼ Exp*(∑_*x*∊ℒ_ *ε*(*x*))
b. Assign *t_min_* = min(*t_imm_, t_frag_, t_exit_*). Assign *T* = *T* + *t_min_*.

- If *t_min_* = *t_frag_*, sample *x_frag_ ∈ L*, such that the probability of sampling *x_i_* is proportional to 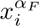. Generate *r ∼ Beta*(*a,* 1). Assign ℒ = ℒ - [*x*_*frag*_] + [*rx*_*frag*_, (1 - *r*)[*x*_*frag*_]
- If *t_min_* = *t_imm_*, generate *x_new_* ∼ *U* (0, *L*). Assign ℒ = ℒ + [*x_new_*].
- If *t_min_* = *t_E_*, sample *x_E_ ∈ L*, such that the probability of sampling *x_i_* is proportional to *ε* (*x_i_*). Assign ℒ = ℒ - [*x_E_*].
c. Assign *ℒ^hist^* = *ℒ^hist^* + ℒ. Assign *d* = *KS_Dist_*(*ℒ, ℒ^hist^*/*T*).

In step 1, the initial conditions of the FRIME process are set up. The list ℒ is the current fragment length distribution and the list *ℒ^hist^* is the list of fragment lengths throughout the history of the simulation.

In the while loop, *κ* is the threshold of tolerance for the metric *d*, which is the Kolmogorov-Smirnov distace between the current fragment list ℒ and the historical average *ℒ^hist^*, where *T* corresponds to the current time step.

In step 2(a), the next immigration, fragmentation and exit times *t_imm_, t_frag_, t_E_* are generated and in step 2(b) the present time *T* is updated to *T* + min{*t_imm_, t_frag_, t_E_*} by the theory of competing exponential. The fragment list *L* is also updated by different mechanisms according to which simulated event comes first.

In step 2(c), the present fragment profile is loaded into the historical fragment list, and the metric *d* is calculated as the Kolmogorov-Smirnov distance between the current fragment list ℒ and the historical average *ℒ^hist^*/*T*. The process terminates when *d* is smaller than some preset tolerance threshold *κ >* 0.

Note that *T* is an auxiliary variable and does not reflect biological time, and is independent of the stationary profile of a FRIME process.

### Computing machines and software

All simulations and clinical data analysis were run on Python 3.9.12 on a MacBook Air M2 2022 with 8GB Memory and the Apple M2 Chip with operating system macOS Ventura 13.3.1. Python packages required for the simulations are Numpy version 1.23.4, Scipy version 1.7.3, Matplotlib version 3.5.1, Sklearn version 1.1.3, Seaborn version 0.11.2 and Pandas version 1.4.2. Simulations of FRIME processes were computationally light; all simulations were run within 3 hours. Clinical data analysis took fewer than 2 hours. All codes for this manuscript are accessible on https://github.com/thltsui/cfDNA-FRIME.

### Data availability and preprocessing

Published cfNano data from Katsman et. al. [44] were used in this study, comprising of cfDNA Nanopore long-read sequencing data from 7 healthy control subjects and 6 lung adenocarcinoma patients. Shallow sequencing was done at about 0.2*×* genome coverage and varies across samples. Anonymised sequence alignment files in BAM format were downloaded from [43].

Alignment and data pre-processing have been previously described in the original publication [44]. Fragment lengths in base pairs were extracted from properly mapped reads using samtools version 1.8 by excluding unmapped reads, non-primary alignment reads, and supplementary alignment reads. Fragment length information was then aggregated to obtain the number of fragments for each unique length in genomic DNA (gDNA) and mitochondrial DNA (mtDNA). We considered gDNA and mtDNA separately because they exhibit very different fragmentation profiles, as the former is nucleosome-bound and has a multi-peak nucleosomal pattern, while the latter is not nucleosome-bound and only has one peak.

### Model fitting and comparison

#### Numerical experiment

To study the effect of each parameter of the FRIME model on its stationary profile, numerical experiments were run where one parameter of interest was assigned multiple values with all other parameters fixed.

For each parameter regime, a stationary profile was generated by running 50 simulations with different seeds. For all experiments, we set *L* = 1, *κ* = 0.05. The sample average, lower quartile and upper quartile of fragment count across samples at each length scale were plotted. The simulated data were also compared to several model-specific best fitting curves for which analytic expressions were available from the theoretical analysis (Table 2). In Section 4 of S1 Appendix, we showed that the evolution of the fragment size distribution of a FRIME process could be described as a solution to a class of partial differential equation. Furthermore, the mean of the stationary distribution of FRIME processes could also be approximated by the stationary solution to the class of partial differential equations. As a result, we could justify that FRIME processes converge to a stationary fragment profile with mean fragment size given by the expression set out in Table 2. For a detailed construction of these stationary best fit curves, we refer to Section 4.2 of the supplementary material.

**Table 2.** Stationary distributions of fragment profiles under different exit mechanisms. For each model, the functions presented describe the expected fragment count per unit length for the stationary distribution of the FRIME process given the extra assumptions on the basic FRIME model. For the CON model, we assumed that the fragmentation ratio was uniformly distributed. For the PFB model, the best fit curve only applies on the region *x > B_E_* and only if the fragmentaiton ratio is uniform. For the PNB model, the stationary distribution curve is applicable if the fragmentation power is *α_F_* = 1. The details of the derivations can be found at Supplementary Section S4.2.

| Model | Extra Assumptions | Stationary solutions ('Best Fit' Curves) |
| --- | --- | --- |
| CON | $r \sim U(0, 1)$ | $C_1(C_F x^{\alpha_F} + C_E)^{-\frac{\alpha_F+2}{\alpha_F}}$ |
| PNB | $\alpha_F = 1$ | $C_3 z^{-\alpha_E} (C_F z^{1-\alpha_E} + 1)^{\frac{3-\alpha_E}{\alpha_E-1}}$ |
| PFB | $r \sim U(0, 1), x > B_E$ | $C_2 x^{-(\alpha_F+2)}$ |

#### FRIME simulation fitting

To test how well FRIME model fits clinical data, stationary profiles from simulated FRIME processes were compared against clinical mitochondrial DNA (mtDNA) fragment profiles from [43].

As fragments longer than 10^4^bp are scarce, and sequencing accuracy for very long fragments is low, all fragments longer than 10^4^ in clinical data were discarded. FRIME processes were simulated with maximum length *L* = 10^4^. Furthermore, since cfDNAs are digested through endonuclease, we took *r ∼ Beta*(1, 1) for our simulations.

Four clinical mitochondrial cell-free DNA fragment profiles sequenced by nanopore technology with more than 500 fragment counts were analyzed in detail. These fragment profiles were compared against simulated fragment profiles from FRIME model under the Kolmogorov-Smirnov test. The simulated fragment profiles were sampled after running a FRIME process under specific parameter regimes after 10^6^ fragmentation / exit/ immigration events.

The genomic cell-free DNA fragment profile for sample ISPRO.bc05 was analyzed, which has the highest sequencing depth among healthy control samples. To adjust for the cyclical fragmentation pattern of genomic DNA (gDNA) data, fragment counts were aggregated according to nucleosomal bins (see S3 Fig for details). The similarity between simulated FRIME profiles and the genomic cfDNA fragment profile was evaluated through a *χ*^2^ goodness of fit test.

#### PFB model fitting

Note that in Table 2, the best-fit curve for PFB model is the simplest among the three options. Furthermore, the PFB model is useful for a broad range of *α_F_* values, while the PNB model is only applicable for *α_F_* = 1. Therefore, a total number of 13 mitochondrial and genomic cell-free DNA fragment profiles in [43] were fitted with the PFB model. Specifically, the tail distributions of these fragment profiles were fitted to a curve of the form *Cx^β^* (Table 2).

## Results

### CfDNA fragment profile data exhibits a peak and a linear decay trend under log-log scale

During initial data exploration, we observed that cell-free mtDNA fragment profile generated from Nanopore long-read sequencing exhibits a linear decay trend under log-log scale. While gDNA fragment profile has a cyclical, nucleosomal pattern, the linear trend remains true for gDNA if fragment counts are averaged over nucleosomal bins. The findings were illustrated using a selected sample in Fig 2. Furthermore, a clear peak in fragment count was observed for all mitochondrial and genomic cfDNA fragment profiles around 100-200bp. Plots for the full set of data can be found in supplementary material S5 Fig S6 Fig S7 Fig.

**Fig 2.**
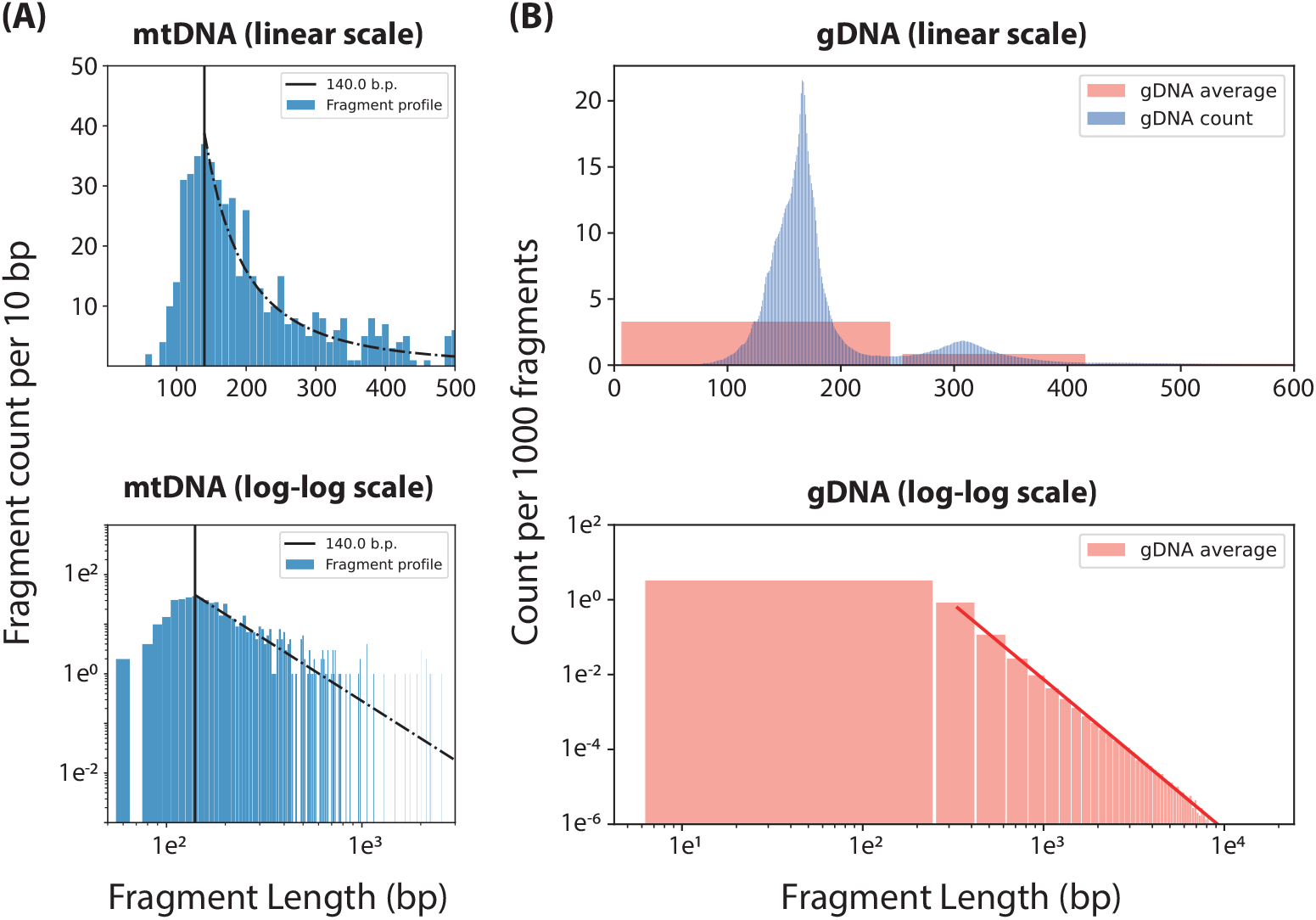
Fragment size profiles of a selected sample ISPRO.bc01 in linear and log-log scale. **(A)** Count per 10bp plotted against mtDNA fragment size shown in linear (top) and log-log scale (bottom). A linear line is fitted against interpolated counts in log-log scale, and is then plotted as a black dashed line in both linear and log-log scale. **(B)** Count per 1000 fragment plotted against gDNA fragment size, shown in linear (top) and log-log scale (bottom). The actual count is shown in blue, whereas the average count for each nucleosomal bin is shown in pink. A best-fit linear line for the average count is plotted in red for log-log scale.

### Exit mechanism determines existence and position of fragmentation profile peak

To study the stationary fragment distribution as a function of fragment size, numerical experiments were run for different parameter values for each of the three classes of exit mechanisms described in Computational model. Some results for specific parameter values are shown in Fig 3.

**Fig 3.**
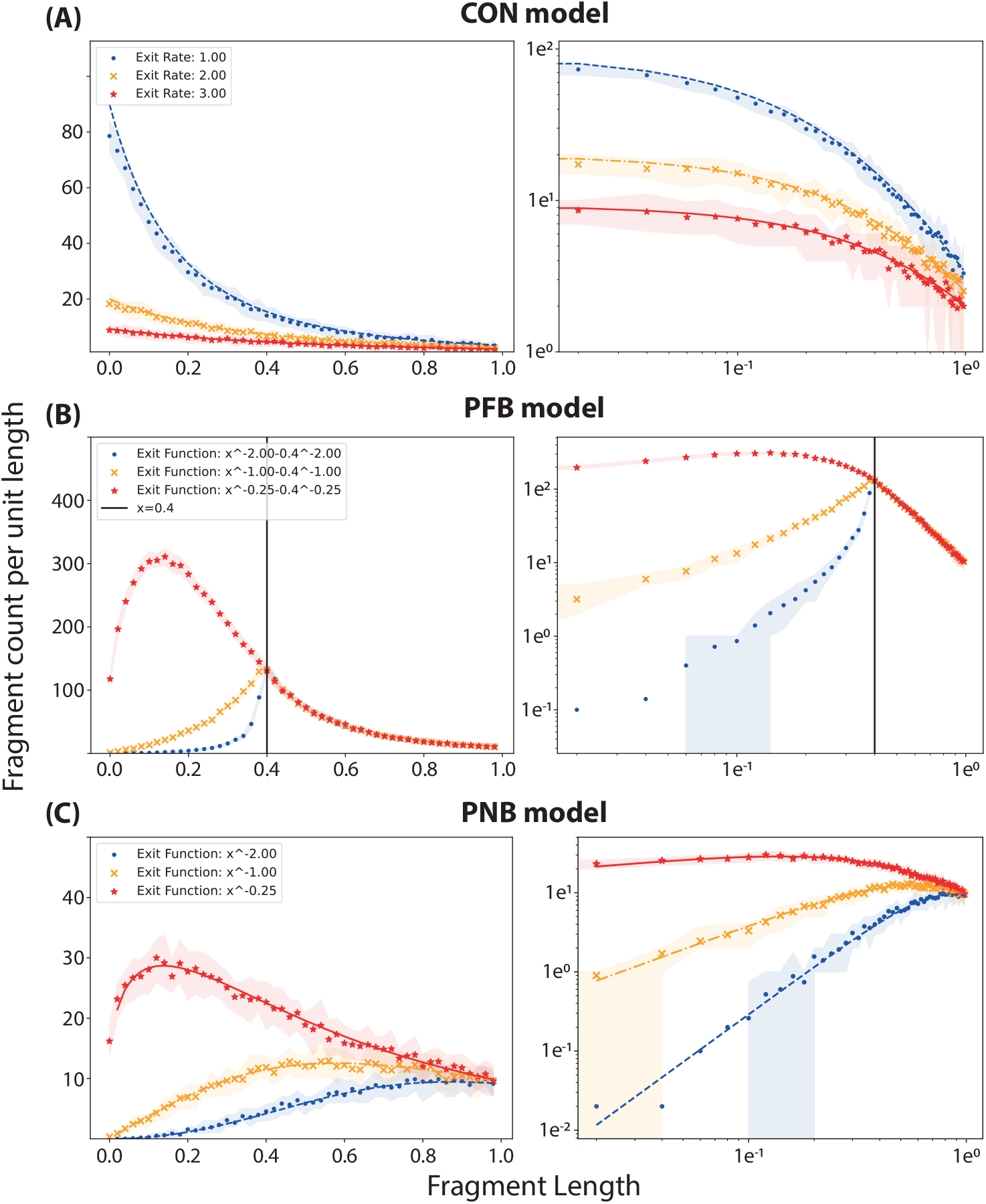
Simulated distribution of fragment sizes for different exit mechanisms. Stationary distributions of fragment sizes obtained from simulations of the FRIME process for parameter values *C_I_* = 500, *C_F_* = 2, *α_F_* = 1, *a* = 1, *κ* = 0.05. 50 points were plotted for each configuration. Each point corresponds to the fragment count per unit length over an interval of length 0.02. Different choices of the exit function were taken and 50 simulations run for each choice of exit function. The plots on the left column are in normal scale, while the plots on the right column are in log-log scale. **(A)** CON model with exit functions *ε* (*x*) = 1 (blue circles), 2 (orange crosses), 3 (red stars). **(B)** PFB model, 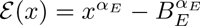, with *α_E_* = -2 (blue circles), -1 (orange crosses), -0.25 (red stars). Parameter *B_E_* was taken to be equal to 0.4. The vertical line was plotted at the exit boundary *B_E_* = 0.4. **(C)** PNB model, *ε* (*x*) = *x^−^*^2^, (blue circles), *x^−^*^1^ (orange crosses), *x^−^*^0.25^ (red stars). Shaded regions were plotted using upper and lower quartile fragment count across all simulations. Best fit lines were also plotted for comparison for the CON and PNB model.

The results in Fig 3 showed relevant trends and features of the different models/exit mechanisms considered. Firstly, the CON model led to a monotonic decrease of the fragment count with the fragment size and did not exhibit a peak in fragment count ubiquitous in clinical cfDNA profiles.

The PFB model showed a peak in the dependence of the fragment count with size. When the boundary played a substantial role, in this case *α_E_ ≤ −*1, the peak was obtained on the boundary *B_E_*, and within the interval [0, *B_E_*] for larger *α_E_* values. For a theoretical justification of the result, refer to Section 4 of S1 Appendix.

Finally, the PNB model displayed a peak number of counts within the interval. The peak moved to smaller fragment counts and was more pronounced as the parameter *α_E_* grew. We note that the log-log plots showed that the tail-distributions for PFB profiles were linearly decreasing in a broad range of small to medium fragment sizes. However, profiles obtained using the CON and PNB models did not exhibit linearity in the log-log plots.

The presence of peaks and the shape of the tail distribution in the log-log plots for the PFB model simulations are consistent with the previous data exploration. Thus, in the following, we perform the subsequent analyses focusing only on the PFB model.

### Fragmentation mechanism determines tail distribution of fragment profile for PFB model

#### Fragmentation ratio and fragmentation power

To study the effect of fragmentation mechanism on stationary fragment profile, numerical experiments were simulated for three different fragmentation ratios and three different fragmentation power rates, as seen in Fig 4.

**Fig 4.**
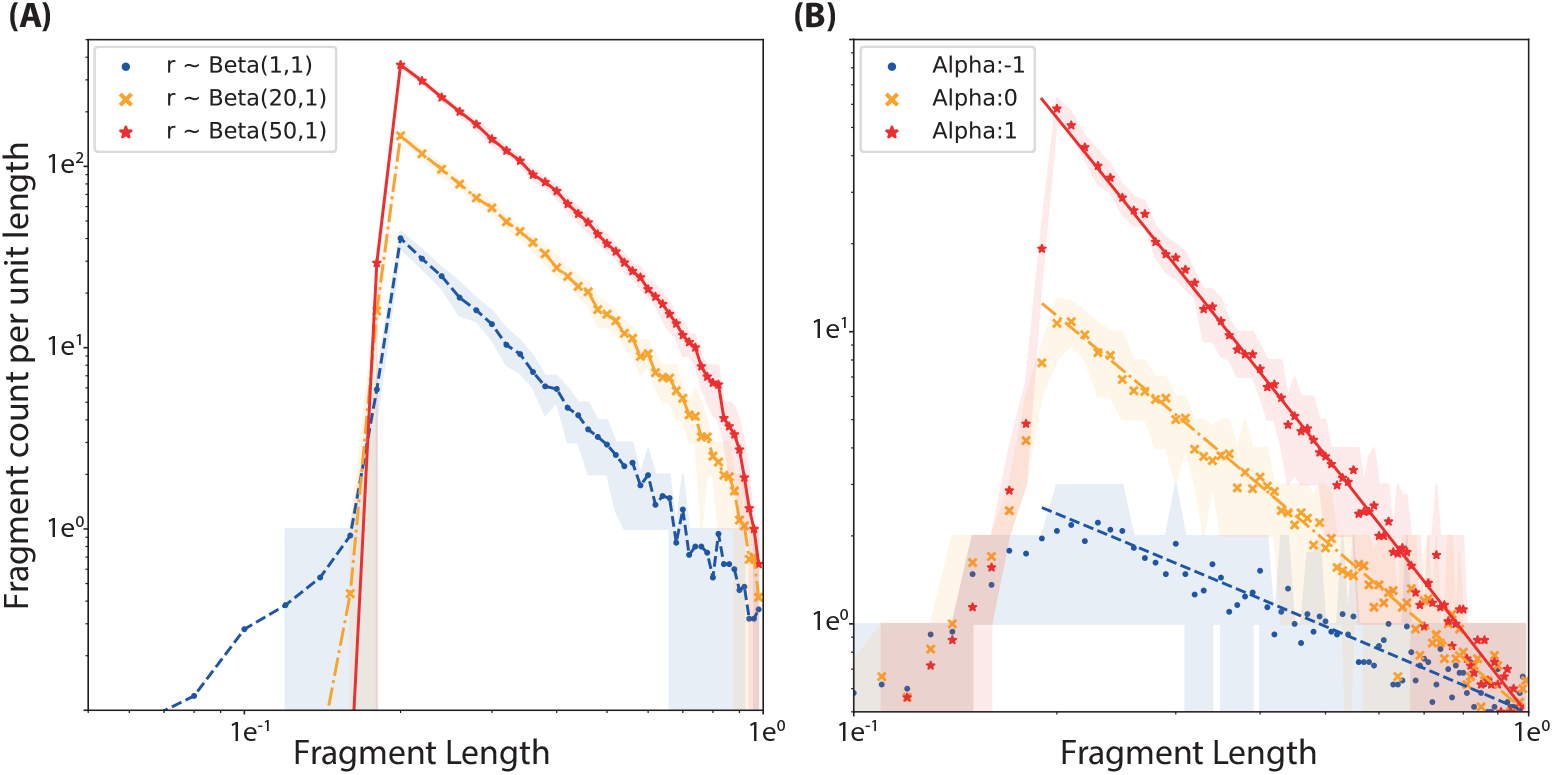
Simulated fragment profiles with different fragmentation ratios and fragmentation power rates. 50 simulations were run for each fragmentation mechanism. 50 points were plotted for each configuration corresponding to average fragment density at intervals sampled in range [0, 1] of length 0.02. Shaded regions were plotted using upper and lower quartile fragment count across all simulations. PFB best fit lines were also plotted for comparison in panel **(B)** with blue-dashed, orange point-dashed and red-solid lines respectively for *α_F_ ∈* {−1, 0, 1}. All plots were in log-log scale. **(A)** The fragmentation ratios were *r ∼ Beta*(1, 1), *Beta*(20, 1), *Beta*(50, 1) respectively for blue-circled, orange-crossed and red-starred dots. Fixed parameters were (*C_I_* = 20, *C_F_* = 1, *α_F_* = 1, *ε* (*x*) = *x^−^*^2^ *−* 0.2^−2^, *κ* = 0.05). **(B)** Fragmentation power were *α_F_ ∈* {−1, 0, 1} respectively for blue-circled, orange-crossed and red-starred dots. Fixed parameters were (*C_I_* = 20, *C_F_* = 1, *r ∼ Beta*(1, 1), *ε*(*x*) = *x*^−2^ − 0.2.^−2^, *κ* = 0.05).

**Fig 5.**
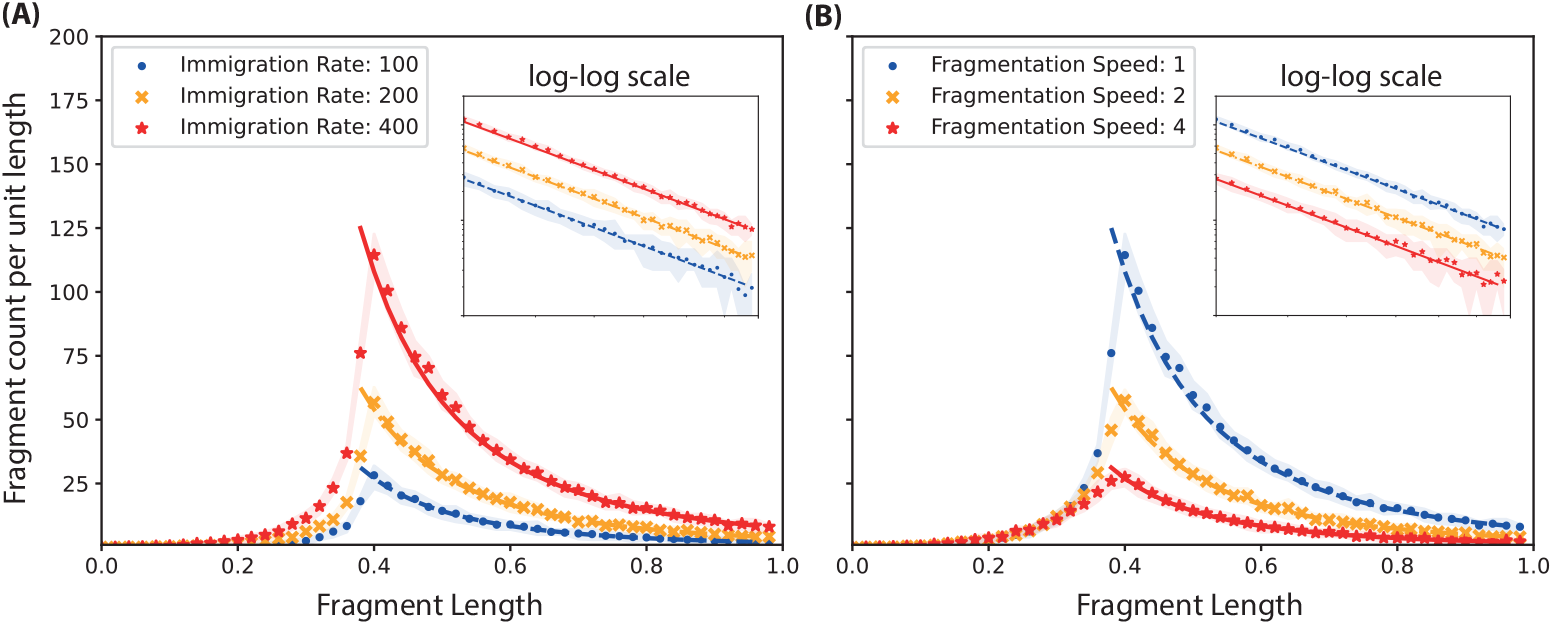
Simulated fragment profiles with different fragmentation and immigration speeds. 50 simulations were run for each fragmentation mechanism. 50 points were plotted for each configuration corresponding to average fragment density at [0,0.02], …[0.98,1]. Shaded regions were plotted using upper and lower quartile fragment count across all simulations. The small plots were the tail distribution but in log-log scale. **(A)** The immigration rates were *C_I_* = 100, 200, 400 for blue circles, orange crosses, and red stars respectively. Fixed parameters were *C_F_* = 1, *α_F_* = 1, *r ∼ Beta*(1, 1), *ε* (*x*) = *x^−^*^2^ *−* 0.4^−2^, *κ* = 0.05. **(B)** The fragmentation speed were *C_f_* = 1, 2, 4 for blue circles, orange crosses, and red stars respectively. Fixed parameters were *C_I_* = 400, *α_F_* = 1, *r ∼ Beta*(1, 1), *ε* (*x*) = *x^−^*^2^ *−* 0.4^−2^, *κ* = 0.05.

In Fig 4(A), several fragmentation profiles with different fragmentation ratio were explored. A larger value in the first coefficient *a* of the *Beta*(*a, b*) distribution (corresponding to exonuclease hypothesis, see Fragmentation mechanisms in Computational model) leads to a steeper decay in the fragment profile tail distribution. In particular when *a* = 1, the fragment profile under the log-log scale is linear.

For fragmentation profile when *r ∼ Beta*(1, 1), the PFB best fit expression of form *Cx^−αF −^*^2^ lies within the shaded region, (see Fig 4(B)). This corresponds to a linear decrease under the log-log scale. Mathematically, if we denote *β_P F B_* to be the slope of the best-fit line in log-log scale, then

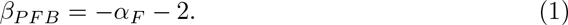

For fragmentation profiles where *r ∼ Beta*(*n, n*) for large *n*, refer to S2 Fig.

#### Fragmentation speed and immigration speed

To study the effect of fragmentation speed on stationary fragment profile, numerical experiments were simulated for three different fragmentation and three different immigration speeds, namely *C_F_* and *C_I_*. The fragmentation speeds were chosen as *C_F_ ∈* {1, 2, 4} with fixed *C_I_* = 400. Similarly, the immigration rates were chosen as *C_I_* = 100, 200, 400 with fixed *C_F_* = 1.

The fragment profiles under the log-log scale are parallel to each other. The fragment profiles are scaled at ratio proportional to the immigration rate and inversely proportional to the fragmentation speed.

For fragment profiles with non-uniform immigration distribution (e.g. Exponential, Normal distributions), refer to S1 Fig.

### Simulated FRIME profiles fit clinical data

Next we wanted to study if the FRIME model was able to describe correctly the different features observed in clinical data. To do so, FRIME processes with different parameters were simulated and compared against the mtDNA fragment profile of the 4 samples ISPRO.S1, ISPRO.bc01, ISPRO.bc03 and ISPRO.bc05 from the nanopore study found in [43]. The 4 samples were chosen according to the criteria described in Model fitting and comparison.

For each of the 4 mtDNA fragment profiles, there are parameter regimes which produce simulated fragment profiles with *p*-values larger than 0.1 under the Kolmogorov-Smirnov test, see Table 3. As the *p*-values are larger than 0.1, the simulated fragment profiles cannot be considered statistically different from clinical data. See Fig 6(A-B) for an example of a simulated FRIME process that fits well with clinical data.

**Fig 6.**
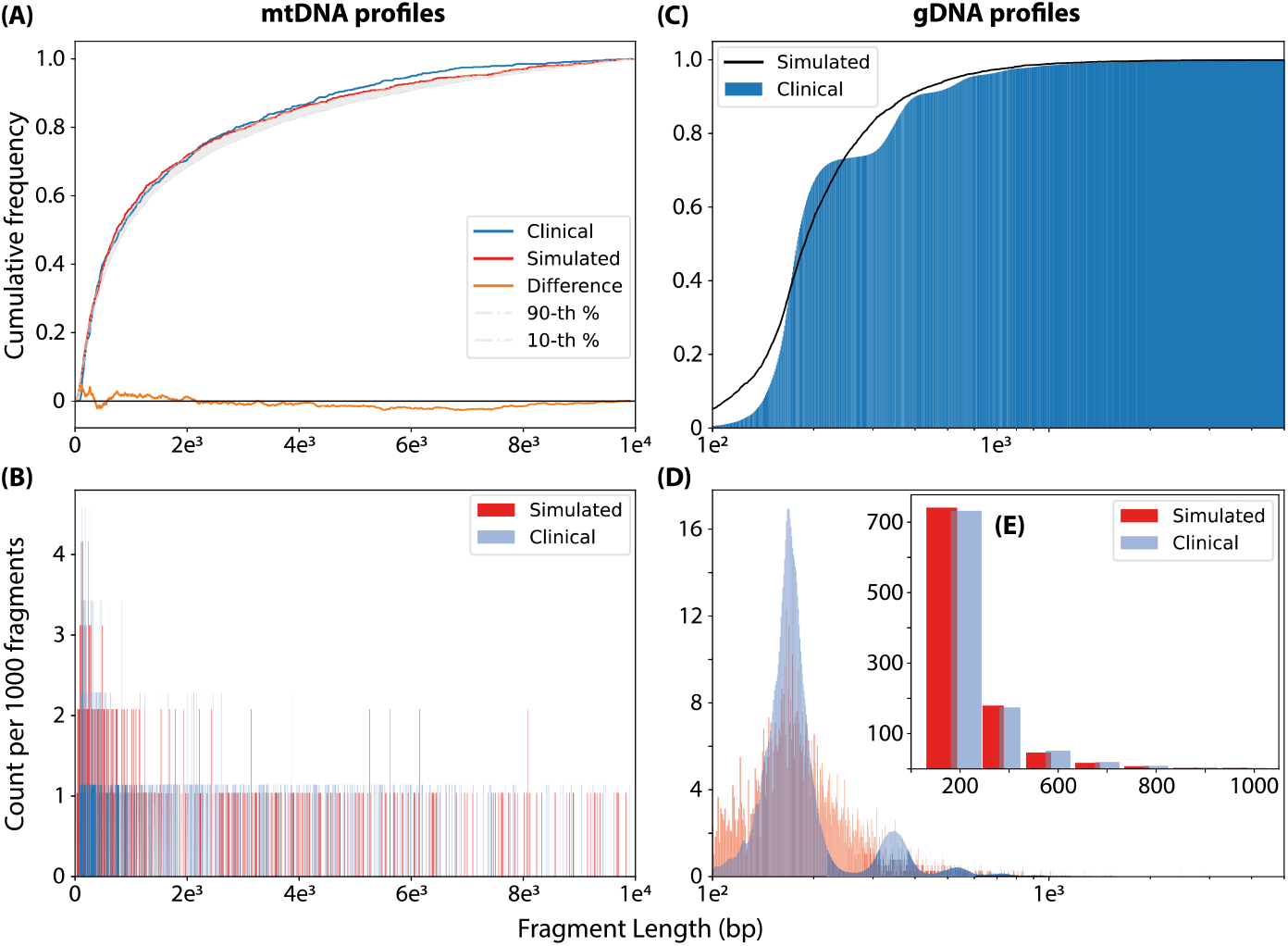
Fitting clinical data for sample ISPRO.bc05 with the FRIME model. **mtDNA -** The mtDNA fragment profile for sample ISPRO.bc05 in nanopore study was compared against a simulated FRIME profile. The simulated profile was generated by running a FRIME model with parameters *C_F_* = 10^6^, *C_I_* = 11, *ε* (*x*) = *L*^2.05^*x^−^*^2.05^, *α_F_* = *−*1, *L* = 10^4^, *r ∼ Beta*(1, 1), with seed = 42 until 10^6^ events occurred. Fragment lengths were rounded to the nearest integer and plotted against clinical data. **(A)** The cumulative fragment count for the mtDNA profile of sample ISPRO.bc05 was plotted as the blue line. The mean cumulative fragment count for the simulated FRIME profile was plotted as the red line, and the shaded region was plotted using 10th and 90th percentile fragment count across all simulations. The difference between the two cumulative count was plotted as the orange line. **(B)** The fragment count (per 1000 fragments) for the mtDNA profile of sample ISPRO.bc05 was plotted as a histogram in blue colour. The fragment count (per 1000 fragments) for the simulated FRIME profile was plotted as a histogram in red colour. Fragment length was measured in base pairs. **gDNA -** The gDNA fragment profile for sample ISPRO.bc05 was compared against a simulated FRIME profile. The simulated profile was generated by running a FRIME model with parameters *C_F_* = 2.5, *C_I_* = 10, *ε* (*x*) = *L*^2^(*x^−^*^2^ *−* 167^−2^), *α_F_* = 1.2, *L* = 10^4^, *r ∼ Beta*(1, 1) until 10^6^ events occurred. Fragment lengths were rounded to the nearest integer and plotted against clinical data. **(C)** The cumulative fragment count for the gDNA profile of sample ISPRO.bc05 was plotted with the blue area. The cumulative fragment count for the simulated FRIME profile was plotted as a black line. The *x*-axis is under the log scale. **(D)** The fragment count for the gDNA profile of sample ISPRO.bc05 was plotted as a histogram in blue colour. The fragment count for the simulated FRIME profile was plotted as a histogram in red colour. Fragment length is measured in base pairs and the *x*-axis is under the log scale. **(E)** The number of fragments in the region [0, 250], [250, 420], [420, 620], [620, 820], *…*, [1220, 1420] for both profiles were plotted as histograms. The simulated profile was plotted in red and the fragment profile for the gDNA profile of sample ISPRO.bc05 was plotted in blue.

**Table 3.** Parameters of mtDNA best fit of the clinical datasets with FRIME processes. Simulated FRIME processes were run for 10^6^ number of events with a fixed initial seed for generating FRIME events. The fragmentation ratio was fixed to be uniform, i.e. *r ∼ Beta*(1, 1), and new fragment lengths were distributed as *U* (0, *L*), with *L* = 10^4^. The column id corresponds to the id of the patient whose mtDNA fragment profile was studied. The constant *C_F_* is the fragmentation speed of the FRIME process and *α_F_* is the fragmentation power. The constant *C_I_* measures the immigration speed. The constant *B_E_* is the exit boundary and *α_E_* the exit power. For PNB model, the exit function is of the form *L^−αE^ x^α_E_^*; for the PFB model, the exit function is of the form 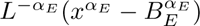. The value *d_KS_* is the Kolmogorov-Smirnov distance between the simulated fragment distribution selected (best fit) and the clinical mtDNA data. The value *p* refers to the *p*-value in the Kolmogorov-Smirnov test.

| id | $C_F$ | $\alpha_F$ | $C_I$ | $\alpha_E$ | Model | $B_E$ | $d_{KS}$ | $p$ |
| --- | --- | --- | --- | --- | --- | --- | --- | --- |
| ISPRO.S1 | $10^6$ | -1.0 | 11.0 | 2.05 | PNB | N/A | 0.022 | 0.287 |
| ISPRO.bc01 | $10^3$ | 0.36 | 10.0 | 2.5 | PFB | 115 | 0.049 | 0.345 |
| ISPRO.bc03 | $10^5$ | -0.5 | 10.0 | 2.10 | PNB | N/A | 0.046 | 0.261 |
| ISPRO.bc05 | $7.5 \times 10^5$ | -0.78 | 8.0 | 2.10 | PNB | N/A | 0.048 | 0.216 |

Similarly, simulated FRIME profiles were found to be statistically close to genomic cfDNA fragment profiles when adjusted for nucleosomal cycles. Under the binning outlined in S3 Fig, simulated FRIME profiles and gDNA profiles were statistically similar for the datasets available. An example is shown in Fig 6(C-E).

### Differences in PFB best-fit lines for gDNA and mtDNA profiles

The tail distribution of gDNA fragmentation profiles for all 13 samples in [43] were fitted with the PFB best-fitting expression of the form *Cx^β^*. Similarly, the mtDNA fragment profiles for 9 samples in [43] were fitted. The PFB best fit lines align with the overall pattern for all of the 9 mtDNA and 13 gDNA fragment profiles as discussed in detail below.

#### Best-fit statistics for mtDNA data

For mtDNA data, the parameter *β_mtDNA_*(slope of the best-fitting line under log-log scale) has an average value of *−*1.56 with range between [*−*2.5, *−*0.69]. According to Equation (1), this corresponds to a negative fragmentation power *α_F_* of around -0.5. See Fig 7 for the best fit expressions of 6 mtDNA profiles. See S5 Fig, S6 Fig for the best-fit curves for all profiles.

**Fig 7.**
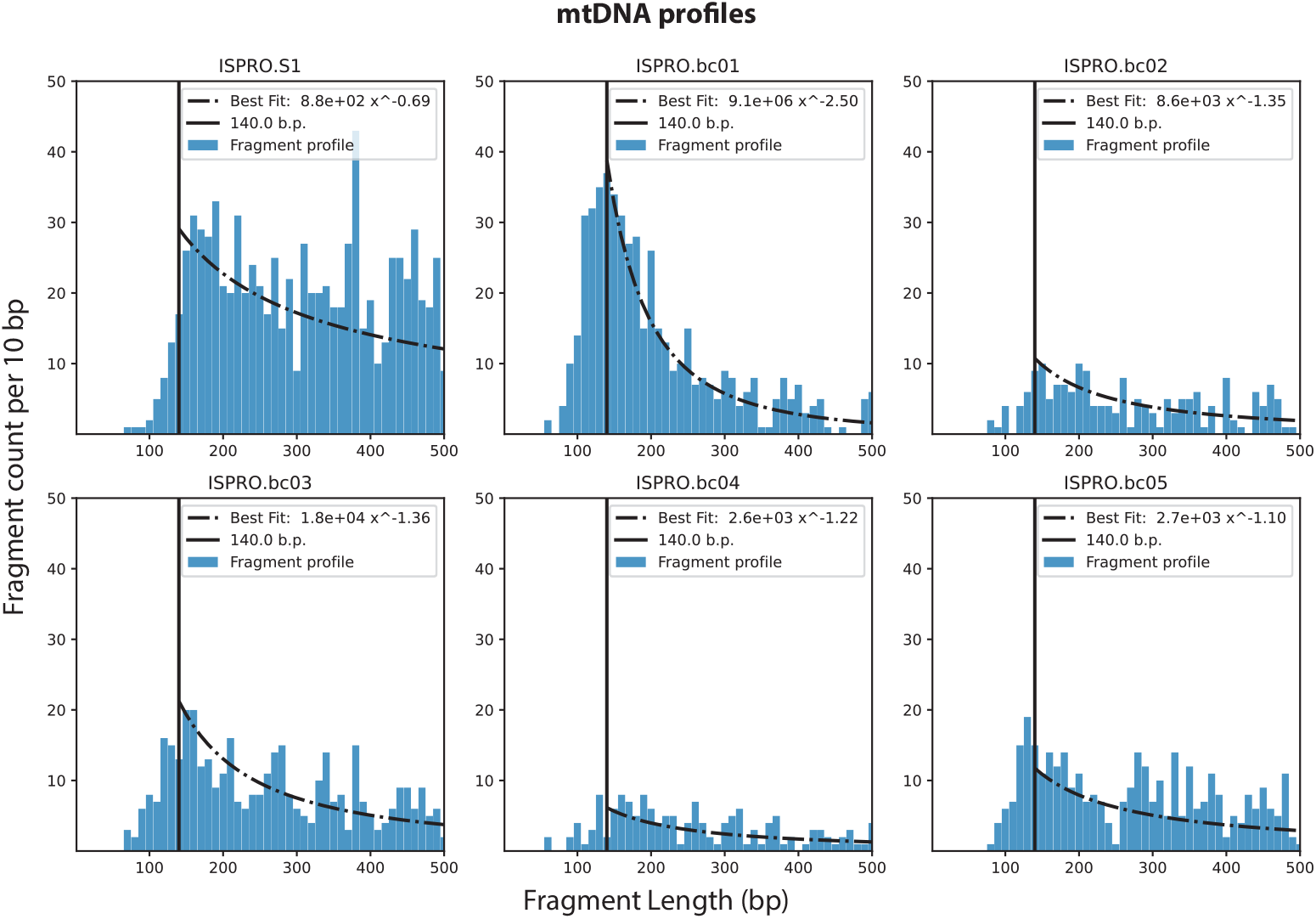
Cell-free mtDNA fragments profiles and PFB best fit lines. Mitochondrial DNA fragment profiles for samples ISPRO.S1, ISPRO.bc01, ISPRO.bc02, ISPRO.bc03, ISPRO.bc04, ISPRO.bc05 from nanopore studies are plotted out as bar charts in blue on the top panels after linear interpolation at intervals of 10bp. Best fit curves of the expression *Cx^β^* are plotted as black dot-dashed lines. The coefficients of the best fit curve are given by a linear fit under the log-log scale on the counts of fragments longer than 140bp. A black vertical line at 140bp indicating the fixed boundary is also plotted on each profile. Both *x* and *y* axes are in linear scale.

#### Best-fit statistics for gDNA data

For gDNA data, the parameter *β_gDNA_* has an average value of *−*3.32 with range between [*−*4, *−*2.5]. According to Equation (1), this corresponds to a positive fragmentation power *α_F_* of around 1. See Fig 8 for the best fit expressions of 6 gDNA profiles. See S7 Fig for the best-fit curves for all profiles.

**Fig 8.**
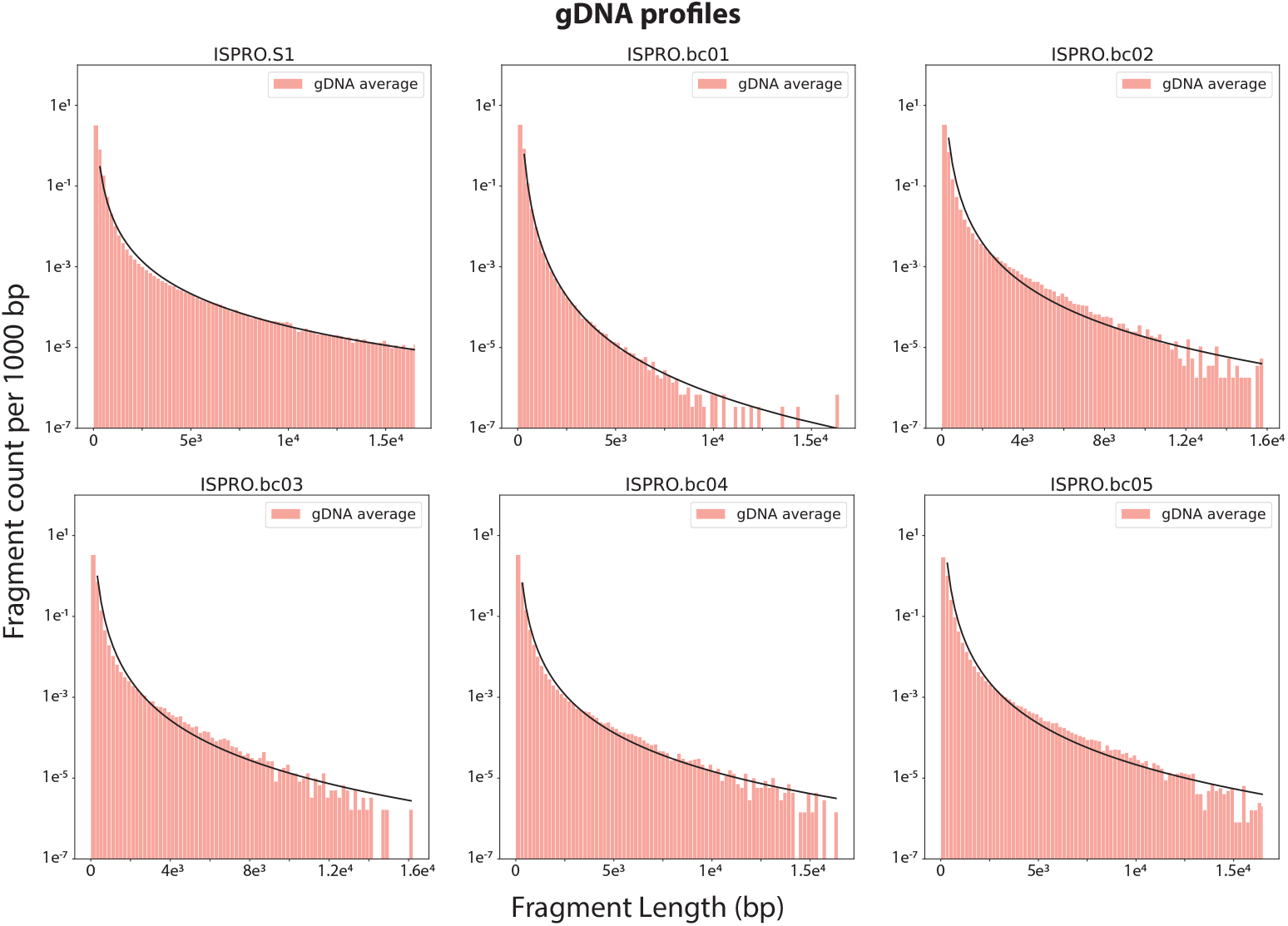
Cell-free gDNA fragment profile with PFB best fit line. Genomic DNA fragment profiles for samples ISPRO.S1, ISPRO.bc01, ISPRO.bc02, ISPRO.bc03, ISPRO.bc04, ISPRO.bc05 from nanopore studies are plotted out as bar charts in pink. Best fit curves of the expression *Cx^β^* are also plotted out. The coefficients of the best fit curve are given by a linear fit under the log-log scale after binning the data. The *y*-axis is in log-scale.

## Discussion

In this paper, we proposed a Markov process (FRIME process) that can model cfDNA production, digestion and clearance in the bloodstream. Our FRIME model is robust and allows us to simulate cfDNA fragment profiles as the equilibrium measure of a FRIME process. Using the FRIME model, we can examine how different assumptions on fragmentation, immigration and exit mechanism produce stationary fragment profiles and compare them against real clinical data. As a result, we can deduce possible cfDNA digestion and clearance mechanisms that are impossible to study under an experimental setting. We can draw the following biological insights from our results.

### Removal mechanism of mitochondrial cell-free DNA is size-dependent and explains the existence of fragment peak

When determining the removal mechanism of cell-free DNA, we considered the cases of size-independent exit (CON), size-dependent exit with no boundary (PNB), and with fixed boundary (PFB) and showed that size-dependent exit mechanism alone is sufficient to result in a peak. On the other hand, size-independent exit results in no peak in the absence of other factors such as protein protection.

Circulating mtDNA, unlike gDNA, is not nucleosome-bound, but demonstrate a single peak at around 100-150bp in the fragment profile, which is variable across individuals. We therefore propose that the existence of a modal fragment size in mitochondrial cfDNA can be explained by its removal mechanism, where shorter fragments are removed faster than longer fragments. Alternatively, the peak can be due to failure to capture ultra-short fragments during sequencing, or protection by protein. However, both cases should result in a uniform peak across individuals, which is not the case.

### Tail distribution of cfDNA fragment profile provides rich information when examined on a log-log scale

In the initial data exploration, we observed that cfDNA fragment profiles exhibit a linear decay trend under log-log scale. In the subsequent modeling of exit mechanism, we showed that only the PFB model has a log-log linear tail distribution, i.e. to the right of the exit boundary. Furthermore, we examined the effect of endonuclease and exonuclease activity on the fragment profile. Log-log linearity is preserved only when all positions have the same chance of being cut (i.e. endonuclease model). In addition, we showed that the slope of the tail distribution is determined by the fragmentation power, which reflects the effect of DNA length on reaction kinetics. Therefore, log-log linearity is a very unique property that can only be met by a few mechanisms, allowing us infer the underlying biological processes.

### Fully nucleosome-bound cell-free DNA is rarely removed from circulation

While there are periodic peaks in genomic cfDNA data, we observed that the fragment distribution binned in nucleosomal units is linear on a log-log scale, until the first nucleosome. From our simulations, this suggests that genomic cfDNA has an exit boundary at around the size of 1 or few nucleosome(s), above which the clearance rate is 0 (corresponding to the PFB model), or very small compared to the fragmentation rate (corresponding to the PNB model with small *α_E_*). We hypothesize that short gDNA, especially those shorter than 167bp, may no longer maintain a stable complex with histone proteins, which leads to a change in physical properties and clearance rate. Our result seems to contradict with previous mice experiments, which have shown that the liver is capable of clearing, or at least trapping, circulating mononucleosomes and chromatin with radiolabelled histones [45, 46]. However, we argue that previous studies only radiolabelled the histone component, and it is unknown whether the nucleosomes are cleared from the circulation as an intact unit with the DNA.

### mtDNA and gDNA undergo different size-biased fragmentation mechanisms

Parameter inference of the tail distribution from clinical data using PFB model showed that mtDNA and gDNA have systematically different slopes, as shown in S8 Fig The fragmentation power *α_F_* can be derived from the slope, and is negative for mtDNA and positive for gDNA. Hence for each molecule, smaller mtDNA fragments react faster than longer mtDNA fragments. Meanwhile, gDNA exhibits the opposite reaction kinetics, and longer fragments react faster than smaller fragments. This suggests that the degradation rate of mtDNA may be diffusion-limited. Smaller fragments react faster because they have a higher diffusion rate than longer ones, which increases the chance of colliding with a DNase. On the other hand, although mtDNA and gDNA are chemically identical, gDNA forms stable complex with histone proteins. This provides an explanation for the positive *α_F_*, which suggests that gDNA fragmentation is limited by surface area instead of diffusion. The longer the DNA molecule, the higher availability of fragmentation sites, hence higher chance of reacting with a DNase. Taken together with the discussion on the clearance mechanism of nucleosome-bound cfDNA, we propose a unifying model where nucleosome-bound cfDNA are not free-floating in circulation, but may have low solubility and form aggregates or phase-separated droplets. This mesoscale organization may sequester the nucleosomes, making them difficult for the body to remove, and cfDNA fragmentation only happens at the contact surface with plasma. This is supported by the existing literature on cell-surface bound DNA, where high molecular weight DNA is found on surfaces of blood cells [50], although the authors did not find low molecular weight DNA by gel electrophoresis, which contradicted with our hypothesis.

### FRIME model allows the interpretation of cfDNA concentration as biomarker of tumor

Plasma cfDNA concentration alone is able to predict patient prognosis in multiple cancer types 47–49 and correlates well with overall tumor burden [47]. Therefore, one hypothesis is that the increased plasma cfDNA comes from increased production. However, literature also claimed that the DNase activity in cancer patients may be reduced, contributing to higher cfDNA concentration [33]. The contributing factors of cfDNA concentration has not been quantitatively studied holistically. From Fig 5, we showed that when the cfDNA fragmentation kinetics remain unchanged, the rate of production of cfDNA is directly proportional to the total amount of fragments in a patient’s bloodstream. This explains the linear correlation between cfDNA concentration and tumor burden. On the other hand, when cfDNA production rate is unchanged, the fragmentation speed is inversely proportional to cfDNA fragment counts. The FRIME model, therefore, provides a numerical framework for understanding the concentration of plasma cfDNA.

### Best fit coefficients in log-log scale can serve as potential biomarkers

We showed in S5 Fig, S6 Fig that all fragment profiles in our study have an excellent fit under the PFB model visually. However, there are variations among individuals on the slope of linearity for such trends. We believe it is possible for this to be a potential biomarker for diseases. Although we do not have enough data to fully examine this hypothesis, preliminary analysis suggests that best-fit gDNA coefficients of patients with lung adenocarcinoma have higher variance than healthy patients. For further detail, refer to S8 Fig.

## Limitation and Future Work

We generated several testable hypotheses from our FRIME model, including the exit mechanisms, physical and chemical kinetic properties for nucleosome-bound and free-floating cfDNA. We hope these ideas can be tested with *in vitro* and *in vivo* experiments in future.

Apart from drawing biological insights, we were also able to infer different model parameters for individual patients, which are related to their cfDNA fragmentation and production rates, which may be more meaningful parameters than cfDNA concentration alone. However, one of the biggest limitations here is the lack of high depth long read cfDNA sequencing data. Most of the published cfDNA sequencing data were done using short-read sequencing platforms, which are very inefficient in capturing DNA molecules beyond 1 kb, thus not suitable for comparing with our model. At the time of our study, only one long read cfDNA sequencing dataset was available without restrictions [43]. Unfortunately, shallow sequencing was done for this dataset. We set an arbitrary cutoff of at least 5000 mtDNA molecules being sequenced in order to statistically compare with our simulation, and only 4 samples passed this cutoff. With more clinical long read sequencing data we will be able to evaluate whether the fitted model parameters from the fragmentation profile are good biomarkers.

Another limitation is that we were not able to reach a definite conclusion about the mode of cfDNA exit mechanism. Although we obtained a good fit using PNB model statistically under the FRIME framework, it cannot be guaranteed that the solution is the best fit.

Finally, our model inferred parameters based on an auxiliary time variable *t*. For example, we may infer *x* fragmentation events per unit time, but further work needs to be done to translate this into the commonly used units for DNase activity. Common used assays of DNase activity involve comparing samples to a reference with known activity [31, 51], which is often specified by the manufacturers as the amount of enzyme required to completely degrade 1 µg of plasmid DNA in the 10 minutes at optimal working conditions. We propose that a simulation can be performed to convert our auxiliary time unit to the degradation assay units. As such, in future studies where the plasma DNase activity is measured alongside with cfDNA sequencing, reasonable fragmentation parameters can be specified prior to fitting our FRIME model, thus allowing a more accurate estimation of the cfDNA production rate.

## Conclusion

We have shown in our clinical data analysis that FRIME model highly recapitulates real life data with a few simple hypotheses, and can be used to draw inferences from mtDNA fragment profile. Although gDNA fragment profile has an inherent cyclical nature, once we adjust our fragment profile by binning data, i.e. by effectively measuring fragment lengths in units of nucleosomes, the gDNA fragment profile also fits into the FRIME model and the log-log linearity model.

We believe that our model reveals insights about cfDNA kinetics that would otherwise be unethical to measure in patients. We look forward to the general application of the FRIME model in other means of liquid biopsy as well, such as circulating RNA and proteins.

## Supporting information

**S1 Appendix Supplementary text for the manuscript** This supplementary test includes further FRIME simulations, a detailed explanation of the statistical test used for clinical data analysis, as well as a mathematical derivation of the best fit curve for CON, PFB, PNB model.

**S1 Fig. FRIME profile with exponential and normal distributed immigration mechanism.** Stationary distributions of fragment sizes obtained from simulations of the FRIME process with immigration and fragmentation parameter values *C_I_*= 500, *C_F_* = 1, *α_F_* = 1, *a* = 1, PFB exit mechanism with parameters *α_E_* = *−*2, *B_E_* = 0.4, and simulation threshold *κ* = 0.05. 50 points were plotted for each configuration. Each point corresponds to the fragment count per unit length over an interval of length 0.02. Different choices of the immigration function were taken and 50 simulations run for each choice of immigration function. Shaded regions were plotted using upper and lower quartile fragment count across all simulations. **(A)** EXP model with immigrating length *z ∼ Exp*(*λ*) where *λ* = 0.1 (dots), 0.5 (*×*), 0.7 (stars). **(B)** The simulated fragment profiles look sufficiently linear under the log-log scale. **(C)** NORM model with immigrating length *z ∼ N* (*µ,* 0.1*µ*) with *µ* = 0.5 (dots), 0.7 (*×*),0.9 (stars). The simulated fragment profiles clearly exhibit a peak at the mean fragment length size.

**S2 Fig. FRIME profile with strong fragmentation around midpoint.** Stationary distributions of fragment sizes obtained from simulations of the FRIME process with uniform immigration and fragmentation parameter values *C_I_* = 100, *C_F_* = 1, *α_F_* = 1, PFB exit mechanism with parameters *α_E_* = *−*2, *B_E_* = 0.2, and simulation threshold *κ* = 0.05. 50 points were plotted for each configuration. Each point corresponds to the fragment count per unit length over an interval of length 0.02. Three different choices of the fragmentation ratios were taken and 50 simulations run for each choice of fragmentation ratio. Shaded regions were plotted using upper and lower quartile fragment count across all simulations. Right panel is normal scale, left panel is log-log scale. **(A)** Fragmentation ratio *r ∼ Beta*(*a, a*) with *a* = 1 (blue circles), 100 (orange crosses), 1000 (red stars). For large *a*, fragments tend to be split into two roughly equal proportions. The simulated fragment profiles exhibit big drops and flat decays. **(B)** In log-log scale, the fragment profiles for large values of *a* look like steps.

**S3 Fig. Genomic DNA data on log-log scale, showing the first 9 partitions.**

**S4 Fig. Evolution of *p*-value under KS test on FRIME simulations.** We ran two FRIME simulations and compared them against the mtDNA fragment profile and gDNA fragment profile for sample ISPRO.bc05 under the two-sample Kolmogorov-Smirnov test. **(A)** A FRIME process with parameters *C_F_* = 7.5 *\** 10^5^, *α_F_* = *−*0.78, *L* = 10^4^, *C_I_* = 8, *ε* (*x*) = *L*^2.10^*x^−^*^2.10^, *r ∼ Beta*(1, 1) is run for 10^5^ events. After every 10^4^ events, the fragment profile of the process is compared against the mtDNA fragment profile under the Kolmogorov-Smirnov test. The *p*-value of the result is plotted in the graph. **(B)** A FRIME process with parameters *C_F_* = 2.5, *α_F_* = 1.2, *L* = 10^4^, *C_I_* = 10, *ε* (*x*) = *L*^2^*x^−^*^2^, *r ∼ Beta*(1, 1) is run for 10^5^ events. After every 10^4^ events, the fragment profile of the process is compared against the gDNA fragment profile under the *χ*^2^ two-sample test. The p-value of the result is plotted in the graph.

**S5 Fig. Mitochondrial cfDNA fragment profiles with best fit curve.** Mitochondrial DNA fragment profiles for all samples from nanopore studies are plotted out as bar charts in blue on the top panels. Best fit curves of the expression *Cx^β^* are plotted as black dot-dashed lines (-.). The coefficients of the best fit curve are given by a linear fit under the log-log scale on the counts of fragments longer than 140bp. A black vertical line at 140bp is also plotted on each profile. The cumulative fragment count for all fragment profiles were plotted out as bar charts in blue. Black lines plotted on top of the bars were linear interpolation of the cumulative fragment count at intervals of 10bp.

**S6 Fig. Mitochondrial cfDNA fragment profiles with best fit curve, log-log scale.** Mitochondrial DNA fragment profiles for all samples from nanopore studies are plotted out. Best fit curve of the expression *Cx^β^* is also plotted out. The coefficients of the best fit curve are given by a linear fit under the log-log scale on the counts of fragments longer than 140bp. A black vertical line at 140bp is also plotted on each profile. Both axes are in log scale.

**S7 Fig. Genomic cfDNA fragment profiles with best fit curve.** Genomic DNA fragment profiles for all samples from nanopore studies are plotted out as bar charts in pink. Best fit curves of the expression *Cx^β^* are also plotted out. The coefficients of the best fit curve are given by a linear fit under the log-log scale after binning the data. The *x*-axis is in log-scale.

**S8 Fig. gDNA best fit coefficients distribution plot.** The best-fit coefficients for linearity in log-log scale were plotted as a scatter plot. The *x*-axis is the gDNA slope, and the *y*-axis is the gDNA intercept. Healthy patients are in blue and patients with lung adenocarcinoma is in pink. The scatter plot suggests that the data for patients with lung adenocarcinoma has higher variability.

**S9 Fig. FRIME profile with different linear exit mechanism.** Stationary distributions of fragment sizes obtained from simulations of the FRIME process with uniform immigration and fragmentation parameter values *C_I_* = 500, *C_F_* = 1, *α_F_* = 1, *a* = 1, *b* = 1 and simulation threshold *κ* = 0.05. 50 points were plotted for each configuration. Each point corresponds to the fragment count per unit length over an interval of length 0.02. Different choices of the immigration function were taken and 50 simulations were run for each choice of immigration function. Shaded regions were plotted using upper and lower quartile fragment count across all simulations. The simulations were run with linear exit mechanism *ε* (*x*) = *B_E_m_E_ − m_E_x*, where *B_E_* = 0.4 and *m_E_* = 2 (blue), 3 (orange), 4 (green). A vertical line at *B_E_* = 0.4 was plotted in black. A best-fit line of 100*x*^3^ was plotted in red in the region *x >* 0.4. Note that when *m_E_* = 4 *>* 3*C_F_*, the peak coincides with 0.4 and when *m_E_* = 2 *<* 3*C_F_*, the peak is to the left of 0.4.

## Acknowledgments

T. H.L. Tsui would like to thank Jere Koskela and Jaromir Sant for their feedback and suggestions. P. F. Xie would like to thank Chi Wu for the interpretation of polymer degradation kinetics and Xin Lu for the encouraging research environment and support. T. H.L. Tsui was partially supported by the EPSRC Centre for Doctoral Training in Mathematics of Random Systems: Analysis, Modelling and Simulation (EP/S023925/1), the Deutsche Forschungsgemeinschaft (DFG, German Research Foundation) under Germany’s Excellence Strategy – EXC-2047/1 – 390685813, St. John’s College Oxford and the Rhodes Trust. P. F. Xie was partially supported by the Croucher Scholarship for Doctoral Study, Ludwig Institute for Cancer Research Oxford branch, Mabel Churn Scholarship, and NDM Prize Studentship. S. Chulían and V. M. Péerez-García are supported by the Spanish Ministerio de Ciencia e Innovacíon, MCIN/AEI/ 10.13039/501100011033 (grant number PID2019-110895RB-I00). V. M. Péerez-García is also supported by Junta de Comunidades de Castilla-La Mancha (SBPLY/21/180501/000145).

## Author contributions

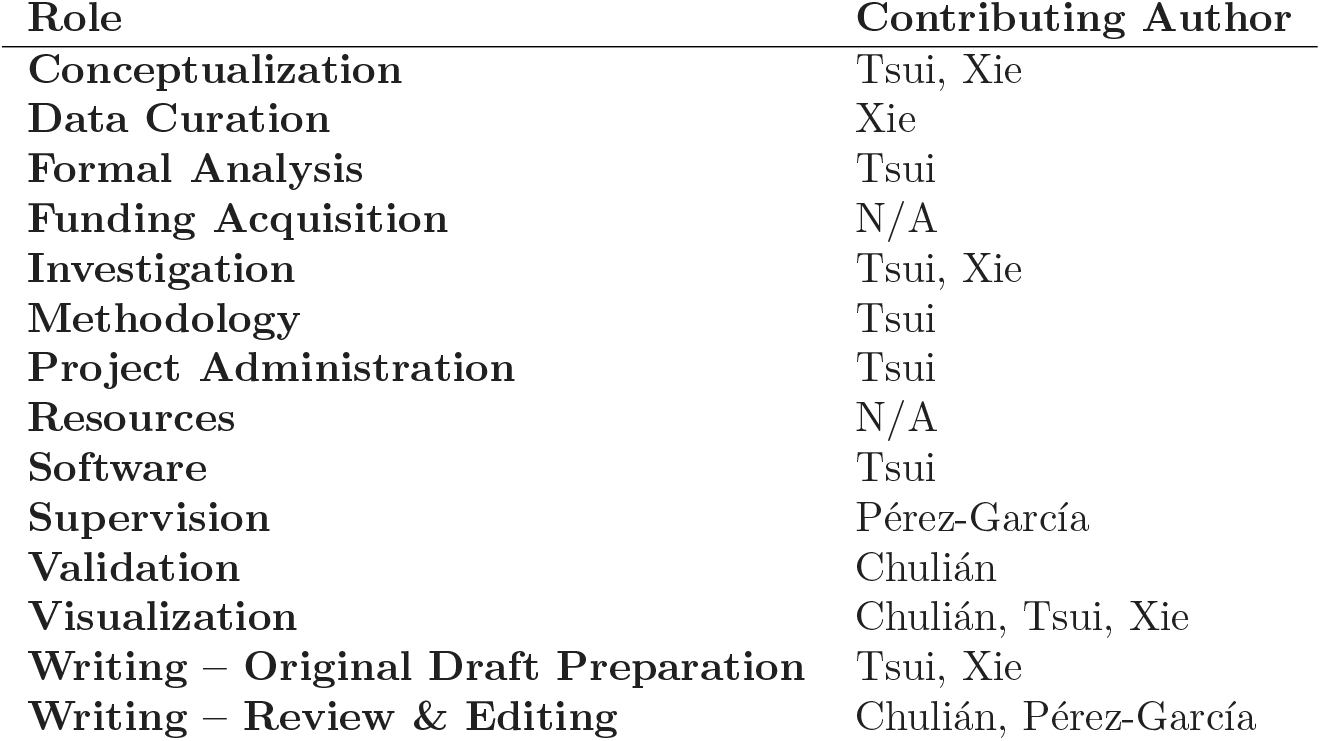

