## Supplementary materials for "Breaking Down Cell-Free DNA Fragmentation: A Markov Model Approach"

July 6, 2023

### 1 Further simulations of FRIME processes

In this section, we include further simulated fragment profiles according to different assumptions on fragmentation, immigration and exit mechanisms.

#### 1.1 Non-uniform immigration mechanism

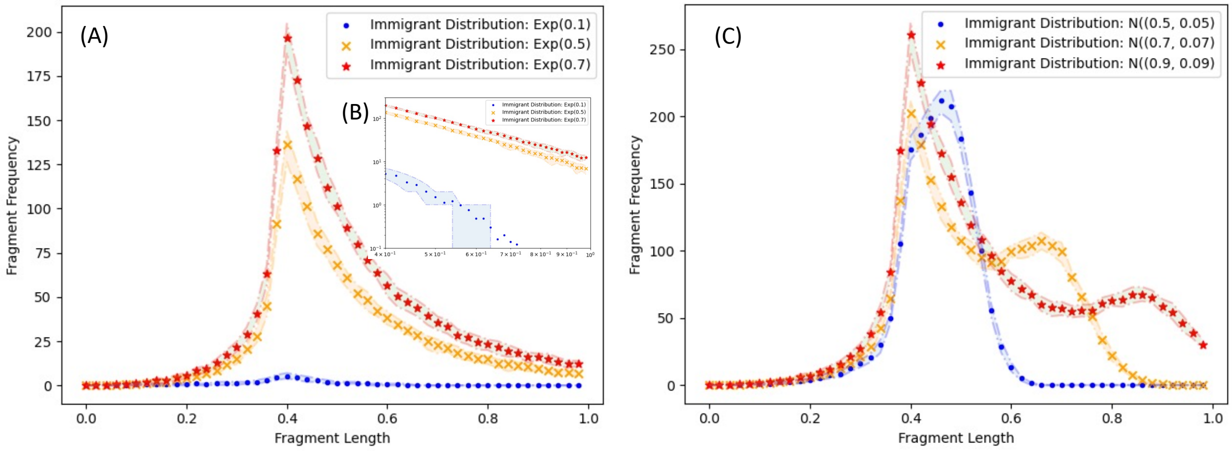


**S1 Fig. FRIME profile with exponential and normal distributed immigration mechanism** Stationary distributions of fragment sizes obtained from simulations of the FRIME process with immigration and fragmentation parameter values $C_I = 500, C_F = 1, \alpha_F = 1, a = 1$, PFB exit mechanism with parameters $\alpha_E = -2, B_E = 0.4$, and simulation threshold $\kappa = 0.05$. 50 points were plotted for each configuration. Each point corresponds to the fragment count per unit length over an interval of length 0.02. Different choices of the immigration function were taken and 50 simulations run for each choice of immigration function. Shaded regions were plotted using upper and lower quartile fragment count across all simulations. **(A)** EXP model with immigrating length $z \sim Exp(\lambda)$ where $\lambda = 0.1$ (dots), 0.5 ($\times$), 0.7 (stars). **(B)** The simulated fragment profiles look sufficiently linear under the log-log scale. **(C)** NORM model with immigrating length $z \sim N(\mu, 0.1\mu)$ with $\mu = 0.5$ (dots), 0.7 ($\times$) ,0.9 (stars). The simulated fragment profiles clearly exhibit a peak at the mean fragment length size.

#### 1.2 Strong fragmentation around mid-point

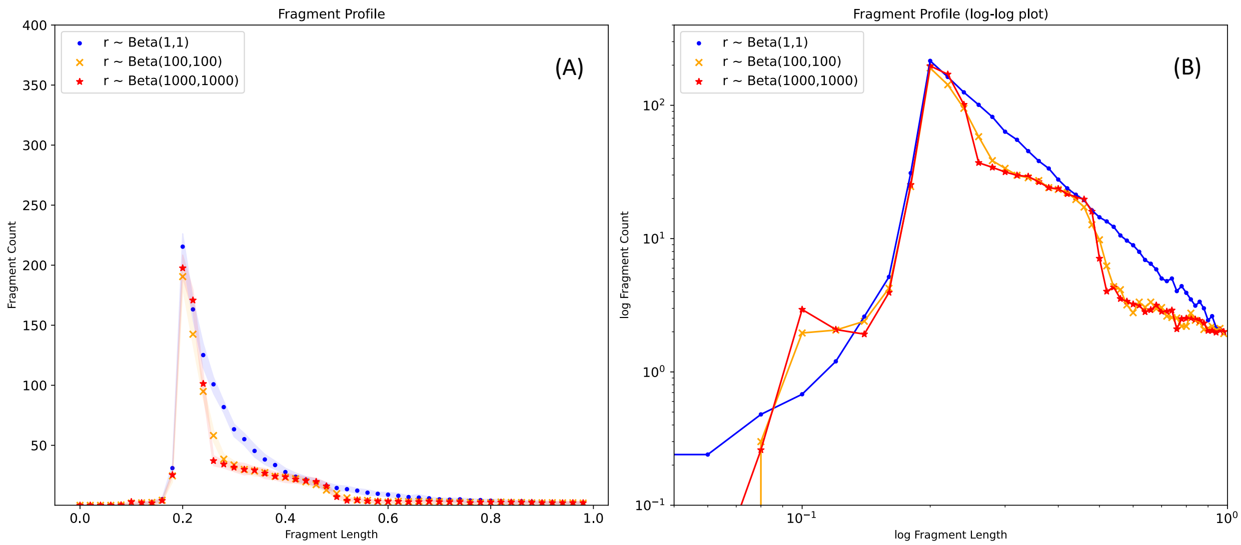


**S2 Fig. FRIME profile with strong fragmentation around mid-point**
Stationary distributions of fragment sizes obtained from simulations of the FRIME process with uniform immigration and fragmentation parameter values $C_I = 100, C_F = 1, \alpha_F = 1$, PFB exit mechanism with parameters $\alpha_E = -2, B_E = 0.2$, and simulation threshold $\kappa = 0.05$. 50 points were plotted for each configuration. Each point corresponds to the fragment count per unit length over an interval of length 0.02. Three different choices of the fragmentation ratios were taken and 50 simulations run for each choice of fragmentation ratio. Shaded regions were plotted using upper and lower quartile fragment count across all simulations. Right panel is normal scale, left panel is log-log scale. **(A)** Fragmentation ratio $r \sim Beta(a, a)$ with $a = 1$ (dots), 100 ($\times$), 1000 (stars). For large $a$, fragments tend to be split into two roughly equal proportions. The simulated fragment profiles exhibit big drops and flat decays. **(B)** In log-log scale, the fragment profiles for large values of $a$ look like steps.

### 2 Clinical data analysis

#### 2.1 Data pre-processing techniques

In this section, we will explain in further detail three techniques employed to process simulated and clinical data for our analysis.

**Nucleosomal binning** To study gDNA data, we binned the fragment counts according to specified nucleosomal bins. This binning is used to account for the cyclical nature of genomic cfDNA fragment profiles.

The nucleosomal bins are defined manually as (0, 250], (250, 420], (420, 620], (620, 820], and every 200bp onward. The binning strategy has been visually confirmed to be reasonable.

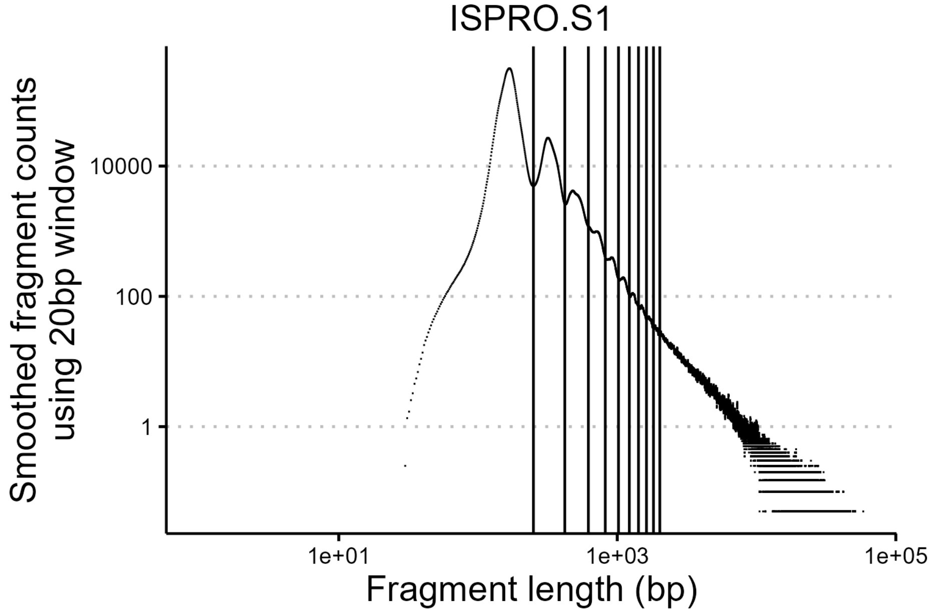


**S3 Fig. Genomic DNA data on log-log scale, showing the first 9 partitions**

**Interpolation method to account for high 0 counts** To fit the PFB best-fitting curve to both mitochondrial and genomic fragment profile, we fitted log fragment count against log fragment length according to a linear model. However, long fragments are sparse and rare in our dataset, so the count of fragments with exactly $x$ base pairs can be 0 if $x$ is large. As a result, under the log scale, the log count becomes negative infinity. To adjust for this sparsity, we plotted the cumulative fragment count against fragment length, then interpolated the count at intervals of 10 bp to give the average fragment count per unit length. We then apply the linear model on the adjusted log fragment counts. Details of the method can be found in the github repository.

**Evolution to stationarity** To evaluate how well the stationary distribution of FRIME processes fit clinical data, we stop a simulated FRIME process after every 10,000 events.

We then conduct a Kolomogorov-Smirnov two-sample test on the simulated fragment profile and the clinical data for mtDNA fragment profile. For gDNA fragment profile, we conduct a $\chi^2$-two-sample test on the total fragment counts in the bins (0, 250], (250, 420], (420, 620], (620, 820].

We observe that the p-values are low initially when the FRIME process has not reached stationarity, and the p-value is consistently above 0.05 after many events. See Figure 4.

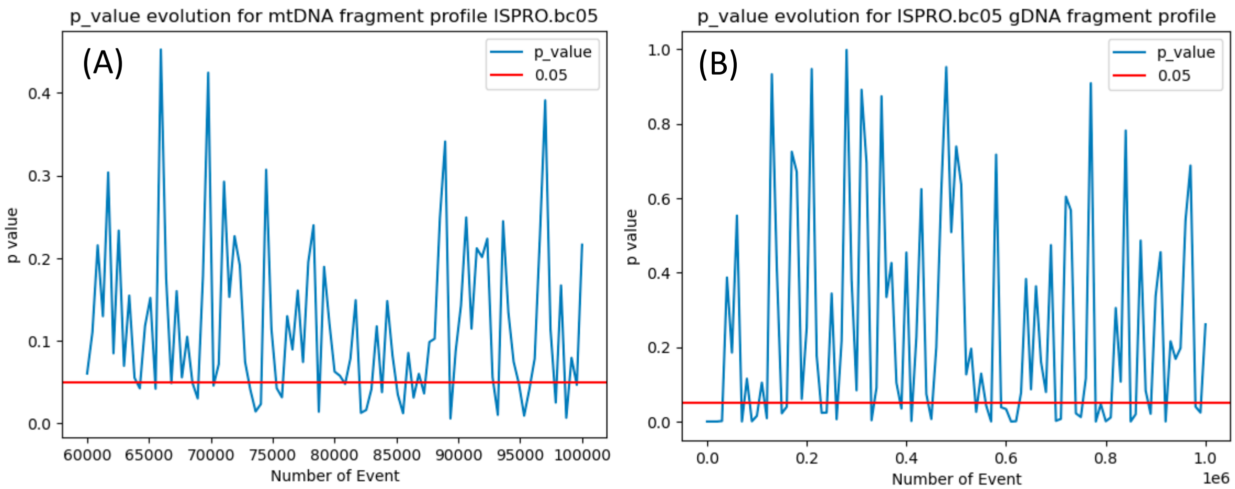


**S4 Fig. Evolution of $p$-value under KS test on FRIME simulations** We ran two FRIME simulations and compared them against the mtDNA fragment profile and gDNA fragment profile for sample ISPRO.bc05 under the two-sample Kolmogorov-Smirnov test. **(A)** A FRIME process with parameters $C_F = 7.5 * 10^5, \alpha_F = -0.78, L = 10^4, C_I = 8, \mathcal{E}(x) = L^{2.10}x^{-2.10}, r \sim Beta(1,1)$ is run for $10^5$ events. After every $10^4$ events, the fragment profile of the process is compared against the mtDNA fragment profile under the Kolmogorov-Smirnov test. The $p$-value of the result is plotted in the graph. **(B)** A FRIME process with parameters $C_F = 2.5, \alpha_F = 1.2, L = 10^4, C_I = 10, \mathcal{E}(x) = L^2x^{-2}, r \sim Beta(1,1)$ is run for $10^5$ events. After every $10^4$ events, the fragment profile of the process is compared against the gDNA fragment profile under the $\chi^2$ two-sample test. The p-value of the result is plotted in the graph.

#### 2.2 PFB best-fitting curves on all samples

In this section we include figures of PFB best-fitting curves on all samples from [1].

##### MtDNA best fit curves

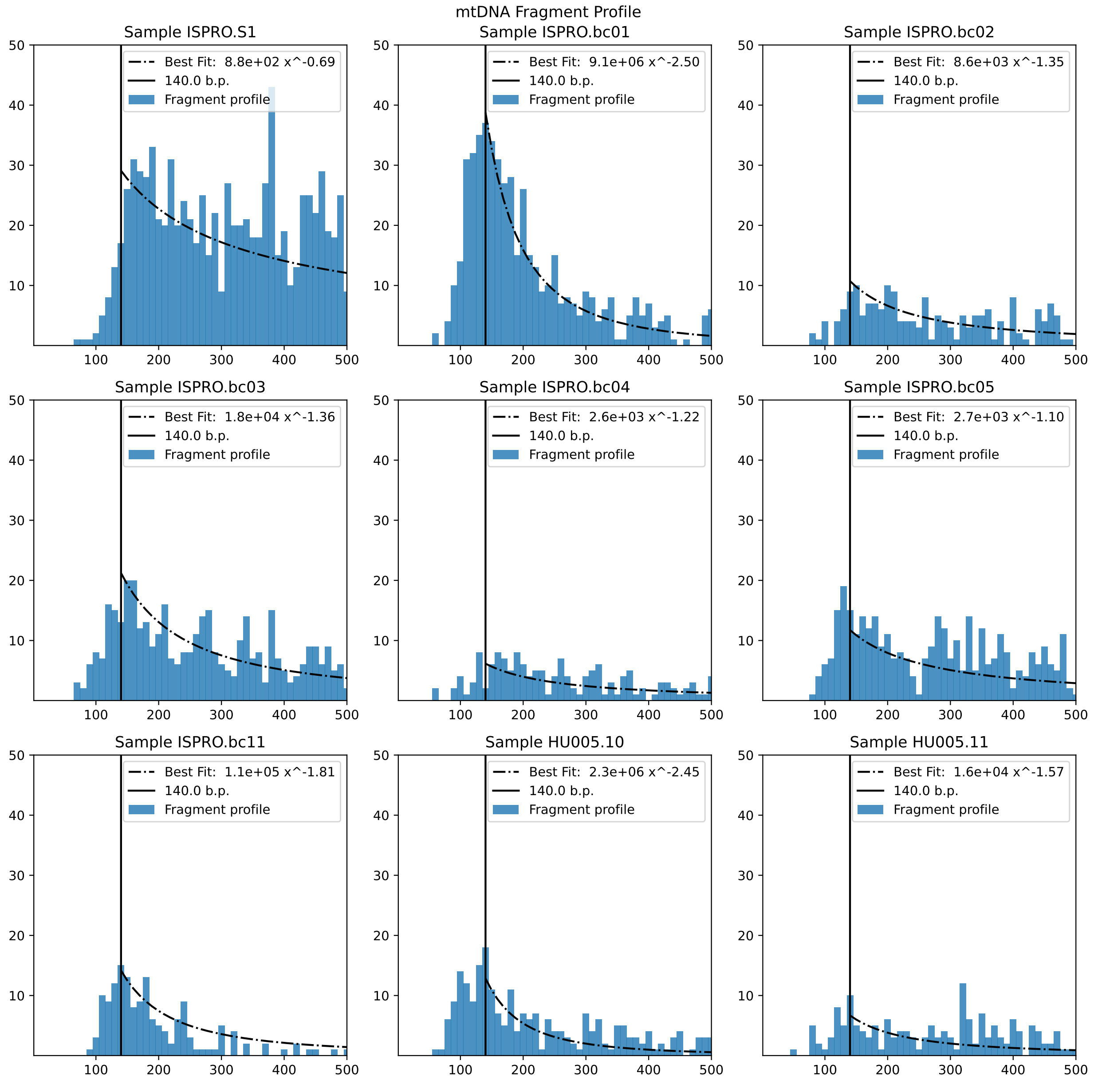


**S5 Fig. mtDNA fragment profiles with best fit curve**

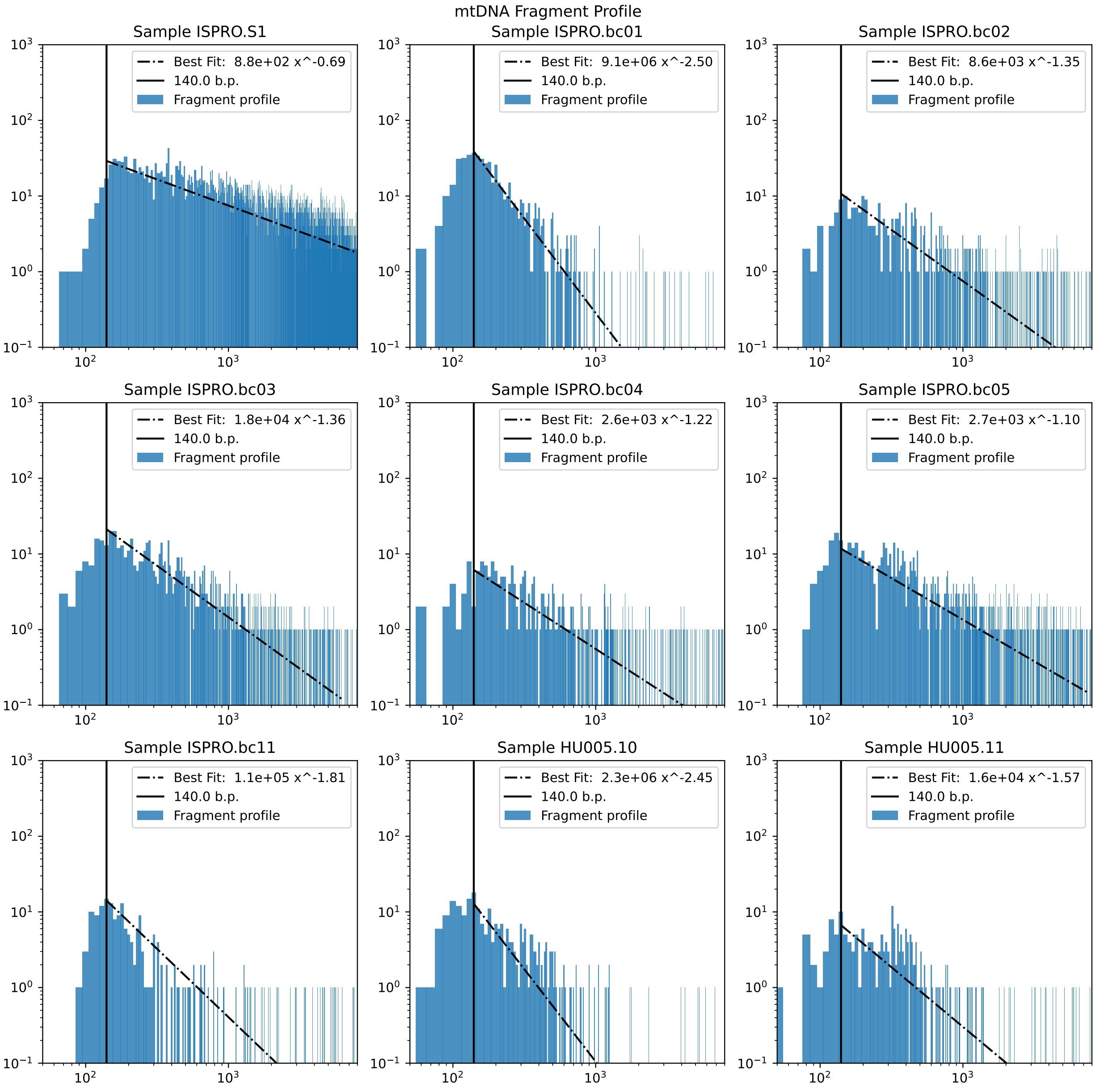


**S6 Fig. mtDNA fragment profiles with best fit curve, log-log scale**

### Genomic DNA best fit curve

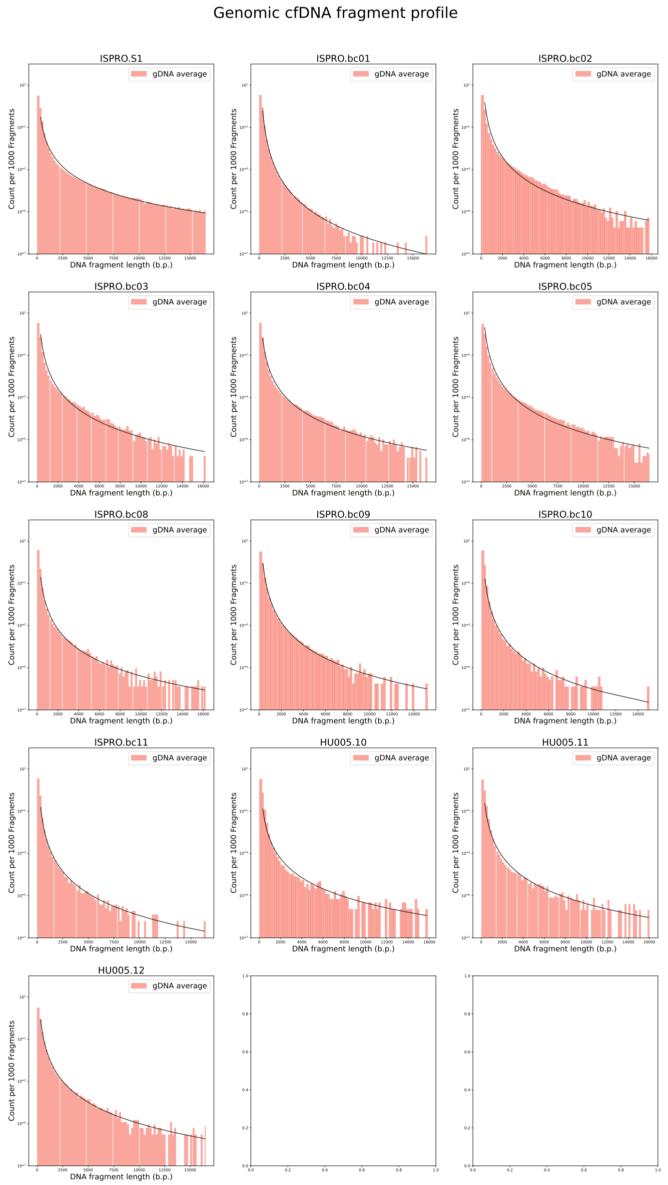


**S7 Fig. Genomic cfDNA fragment profiles with best fit curve**

**Distribution of gDNA and mtDNA best fit coefficients**

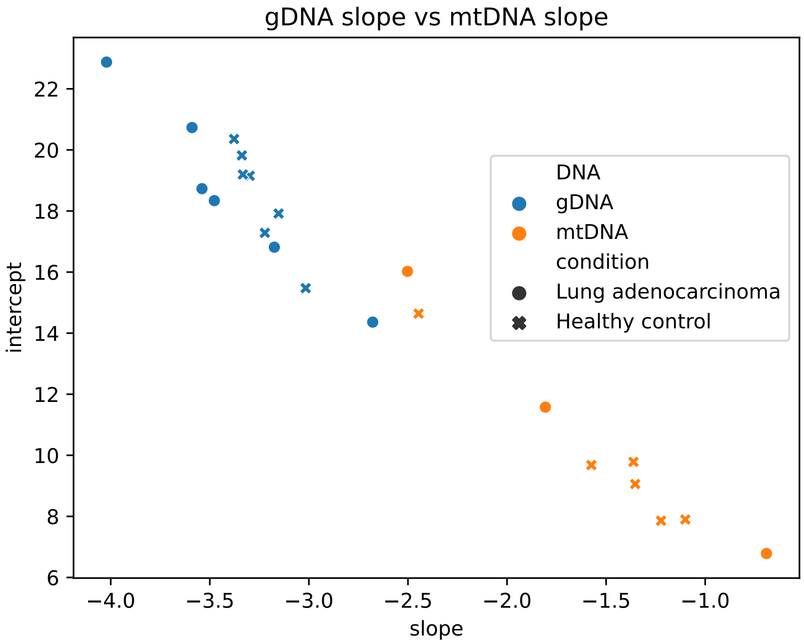


**S8 Fig. gDNA and mtDNA best fit coefficients distribution plot**

### 3 Sample Mean and Hydrodynamic Limit of FRIME Processes

In this section, we will apply the law of large numbers to equate the sample mean of a FRIME process with its hydrodynamic limit. We will also show that the hydrodynamic limit is given by a *fragmentation with immigration and exit* equation.

#### 3.1 Definition of FRIME processes

**Fragmentation with exit processes** First we call a Markov process $(F_X(t))_{t\geq 0}$ a $(C_F, \alpha_F, a, b, \ell, L, \mathcal{E}(x))$-fragmentation with exit process if it evolves according to the following dynamics.

- The process starts off with a fragment with length $x_i(0) \sim U(0, L)$.
- Each fragment of length $x$, at rate $C_F x^{\alpha_F}$, fragments into two pieces of length $xr$ and $x(1-r)$, where $r \sim Beta(a, b)$.
- Each fragment of length $x$, at rate $\mathcal{E}(x)$, leaves the system. All fragments with size smaller than $\ell$ exit the system instantaneously.

In particular, we only consider three types of exit function $\mathcal{E}(x)$, the constant (CON) model, the power exit with fixed boundary (PFB) model and the power exit with no boundary (PNB) model. For the CON model, $\mathcal{E}(x) = C_E$. For the PNB model, $\mathcal{E}(x) = L^{-\alpha_E} x^{\alpha_E}$, $\alpha_E < 0$. FOr the PFB model, $\mathcal{E}(x) = L^{-\alpha_E} \max\{x^{\alpha_E} - B_E^{\alpha_E}, 0\}$.

We denote the empirical measure of fragment lengths present in the system at time $t$ by $F_X(t) := \sum_{i=1}^{N(t)} \delta_{x_i}$, where $x_1, ..., x_{N(t)}$ are the fragments present at time $t$. We also denote the set of continuous functions on the interval $[\ell, L]$ as $C(\ell, L)$.

Finally, for any function $f \in C([\ell, L])$, we define

$$\langle f, F_X(t) \rangle := \sum_{i=1}^{N(t)} f(x_i) \tag{3.1}$$

to be the sum of $f(x)$ over all fragment lengths at time $t$.

**FRIME processes** Similarly, we call $(F_{IX}(t))_{t\geq 0}$ a $(C_F, \alpha_F, a, b, C_I, \ell, L, \mathcal{E}(x))$-FRIME process if it evolves according to the following dynamics.

- The process starts off with a fragment with length $x_i(0) \sim Uni[0, L]$.
- Fragments of length $x$ at rate $C_F x^{\alpha_F}$ fragments into two pieces of length $xr$ and $x(1-r)$, where $r \sim Beta(a, b)$.
- Fragments of length $x$ leave the system at rate $\mathcal{E}(x)$.
- New fragments immigrate into the system according to a Poisson process with intensity $C_I$. The length $z$ of a new fragment is given by the distribution with probability density $p_I(s)ds$.

In other words, the FRIME process $(F_{IX}(t))_{t\geq 0}$ is a $F_X$-with-immigration process defined in [2].

#### 3.2 Law of large number on FRIME processes

We now consider $n$ independent and identically distributed FRIME processes $(F_{IX}^{(i)}(t))_{t\geq 0}$, $i = 1, .., n$. For all continuous functions $f \in C([\ell, L])$ on the interval $[\ell, L]$, we define the random variable

$$\langle f, F_{IX}^{(i)}(t) \rangle := \sum_{x \in F_{IX}^{(i)}(t)} f(x) \tag{3.2}$$

to be the sum of $f$ over lengths of all fragments at time $t$ for the i-th FRIME process.

Note that the random variables $\langle f, F_{IX}^{(i)}(t)\rangle,\ i = 1, ..., n$ are real-valued, independent and identically distributed. Therefore, by the Strong Law of Large Number, the empirical average of $n$ i.i.d samples of FRIME processes will converge to the expected value of another related FRIME process, i.e.

$$\lim_{n\to\infty} \frac{1}{n}\sum_{i=1}^{n} \langle f, F_{IX}^{(i)}(t)\rangle = \mathbb{E}\langle f, F_{IX}^{(1)}(t)\rangle. \tag{3.3}$$

Therefore, to understand the mean of the FRIME process, it suffices for us to understand the limit of $\frac{1}{n}\sum_{i=1}^{n} F_{IX}^{(i)}(t)$ under the weak topology.

Since individual fragments of FRIME processes evolve independently of each other, the FRIME processes satisfy the superposition property, i.e. for two FRIME processes satisfying $F_{IX}^{\mu}(t) = \mu, F_{IX}^{\nu}(t) = \nu$ at time $t$, for all $s > 0$

$$F_{IX}^{\mu}(t+s) + F_{IX}^{\nu}(t+s) =^d F_{IX}^{\mu+\nu}(t+s), \tag{3.4}$$

where $(F_{IX}^{\mu+\nu}(t+s))_{s\geq 0}$ is an independent FRIME process that satisfies $F_{IX}^{\mu+\nu}(t) = \mu + \nu$.

Furthermore, the sum of $n$ independent Poisson point processes of intensity $C_I$ has the same distribution as a Poisson point process of intensity $nC_I$. Therefore, the sum of $n$ i.i.d $(C_F, \alpha, a, b, C_I, \ell, L, \mathcal{E}(x))$-FRIME process is again a FRIME process, albeit with a different immigration rate. Explicitly, the process is a $(C_F, \alpha, a, b, nC_I, \ell, L, \mathcal{E}(x))$-FRIME process, with all but one parameters being identical to those of the original Frime process, and a new immigration rate of $nC_I$.

As a result, the mean of a FRIME process is the hydrodynamic limit when we scale intensity of immigration by a factor of $n$, and the mass of a fragment by a factor of $1/n$.

#### 3.3 Hydrodynamic Limit of FRIME Processes

**Prelimit Model:** For a $(C_F, \alpha, a, b, C_I, \ell, L, \mathcal{E}(x))$-FRIME process $F_{IX}(t)$, we write $F_{IX}^n(t)$ to be the associated $(C_F, \alpha, a, b, nC_I, \ell, L, \mathcal{E}(x))$-FRIME process. Similarly, we write $\mu_t^n$ to be the re-scaled empirical distribution of fragment lengths at time $t$ such that each fragment has a mass of $1/n$, i.e.

$$\mu_t^n := \frac{1}{n}\sum_{x\in F_{IX}^n(t)} \delta_x.$$

For each $f \in C([\ell, L])$, we define $X_t^{n,f}$ to be the real-valued projection of $\mu_t^n$ through $f$, i.e.

$$X_t^{n,f} := \frac{1}{n}\sum_{x_i\in F_{IX}^n(t)} f(x_i) = \int f(x)\mu_t^n(dx). \tag{3.5}$$

**Identifying the Limit:** We evaluate the evolution of $X_t^{n,f} := \int f(x)\mu_t^n(dx)$ for all $f \in \mathcal{C}([\ell, L])$.

To simplify our calculation, we use the notation $\mathbb{E}_r[g(r)] = \int_0^1 g(r)\frac{\Gamma(a+b)}{\Gamma(a)\Gamma(b)} r^{a-1}(1-r)^{b-1}dr$ to denote the mean of a function $g(r)$ under the $Beta(a,b)$ distribution.

$$\begin{aligned}
&\lim_{\delta t\to 0}\mathbb{E}\left[\frac{X^{n,f}_{t+\delta t}-X^{n,f}_t}{\delta t}\middle|\mu^n_t=\nu\right]\\
=&\int_\ell^L C_F x^{\alpha_F}\mathbb{E}_r\left[\frac{f(xr)\mathbb{1}_{xr\geq\ell}+f(x(1-r))\mathbb{1}_{x(1-r)\geq\ell}-f(x)}{n}\right]\times n\nu(dx)\\
&+nC_I\int_\ell^L\frac{f(x)}{n}p_I(x)dx-\int_\ell^L\mathcal{E}(x)\frac{f(x)}{n}\times n\nu(dx)\\
=&\int_\ell^L C_F x^{\alpha_F}\mathbb{E}_r\left[f(xr)\mathbb{1}_{xr\geq\ell}+f(x(1-r))\mathbb{1}_{x(1-r)\geq\ell}-f(x)\right]\nu(dx)\\
&+C_I\int_\ell^L f(x)p_I(x)dx-\int_\ell^L\mathcal{E}(x)f(x)\nu(dx).
\end{aligned}\tag{3.6}$$

Similarly,

$$\begin{aligned}
&\lim_{\delta t\to 0}\mathbb{E}\left[\frac{(X^{n,f}_{t+\delta t}-X^{n,f}_t)^2}{\delta t}\middle|\mu^n_t=\nu\right]\\
=&\int_\ell^L C_F x^{\alpha_F}\mathbb{E}_r\left[\frac{(f(xr)\mathbb{1}_{xr\geq\ell}+f(x(1-r))\mathbb{1}_{x(1-r)\geq\ell}-f(x))^2}{n^2}\right]\times n\nu(dx)\\
&+nC_I\int_\ell^L\frac{f^2(x)}{n^2}p_I(x)dx+\int_\ell^L\mathcal{E}(x)\frac{f^2(x)}{n^2}\times n\nu(dx)\\
=&\frac{1}{n}\int_\ell^L C_F x^{\alpha_F}\mathbb{E}_r\left[(f(xr)\mathbb{1}_{xr\geq\ell}+f(x(1-r))\mathbb{1}_{x(1-r)\geq\ell}-f(x))^2\right]\nu(dx)\\
&+\frac{1}{n}C_I\int_\ell^L f^2(x)p_I(x)dx+\frac{1}{n}\int_\ell^L\mathcal{E}(x)f^2(x)\nu(dx)
\end{aligned}\tag{3.7}$$

In other words, $(\mu^n_t)_{t\geq 0}$ is a solution to the following martingale problem:
For all $f\in C(\ell,L)$,

$$\begin{aligned}
M^{n,f}_t:=&\int f(x)\mu^n_t(dx)-\int f(x)\mu^n_0(dx)\\
&-\int_0^t\Bigg\{\int_\ell^L C_F x^{\alpha_F}\mathbb{E}_r\left[f(xr)\mathbb{1}_{xr\geq\ell}+f(x(1-r))\mathbb{1}_{x(1-r)\geq\ell}-f(x)\right]\mu^n_s(dx)\\
&\qquad+C_I\int_\ell^L f(x)p_I(x)dx-\int_\ell^L\mathcal{E}(x)f(x)\mu^n_s(dx)\Bigg\}ds
\end{aligned}\tag{3.8}$$

is a martingale, with quadratic variation

$$\begin{aligned}
[M^{n,f}]_t=&\frac{1}{n}\int_0^t\Bigg\{\int_\ell^L C_F x^{\alpha_F}\mathbb{E}_r\left[(f(xr)\mathbb{1}_{xr\geq\ell}+f(x(1-r))\mathbb{1}_{x(1-r)\geq\ell}-f(x))^2\right]\mu^n_s(dx)\\
&\qquad+C_I\int_\ell^L f^2(x)p_I(x)dx+\int_\ell^L\mathcal{E}(x)f^2(x)\mu^n_s(dx)\Bigg\}ds.
\end{aligned}\tag{3.9}$$

As $n\to\infty$, the quadratic variation term $[M^{n,f}]_t\to 0$. Assuming tightness, we have $M^{n,f}_t\to 0$. As a result, the real-valued process $(X^{n,f}_t)_{t\geq 0}$ converges in the Skorokhod topology to a deterministic process

$(X_t^{\infty,f})_{t\geq 0} := (\int f(x)\mu_t^\infty(dx))_{t\geq 0}$, which solves the FRIME equation

$$\begin{aligned}\frac{\partial}{\partial t}\int f(x),\mu_t^\infty(dx) =& C_F\int_\ell^L x^{\alpha_F}\mathbb{E}_r\left[f(xr)\mathbb{1}_{xr\geq\ell}+f(x(1-r))\mathbb{1}_{x(1-r)\geq\ell}-f(x)\right]\mu_t^\infty(dx)\\ &-\int_\ell^L \mathcal{E}(x)f(x)\mu_t^\infty(dx)+C_I\int f(z)p_I(z)dz,\end{aligned} \tag{3.10}$$

with initial condition $\mu_0^\infty(dx)=\mathbb{1}_{[\ell,L]}(x)\frac{1}{L-\ell}dx$.

### 4 Solutions to the FRIME equations

We now consider more explicit expressions for solutions to the FRIME equation (3.10).

#### 4.1 Density solution to FRIME equations

We assume that the solution $\mu_t^\infty(dx)$ is differentiable in space and time, i.e. there exists a function $u_t(x)$ : $[0,\infty)\times[\ell,L]\mapsto\mathbb{R}$ differentiable in $t$ and continuous in $x$ such that $\mu_t^\infty(dx)=u_t^\infty(x)dx$.

Furthermore, recall that for $r\sim Beta(a,b)$, the ratio $\tilde{r}:=1-r$ is distributed as $Beta(b,a)$. For simplicity, we once again use the notation $\mathbb{E}_r[g(r)]=\int_0^1 g(r)\frac{\Gamma(a+b)}{\Gamma(a)\Gamma(b)}r^{a-1}(1-r)^{b-1}dr$ and similarly, we write

$$\mathbb{E}_{\tilde{r}}[g(\tilde{r})]=\int_0^1 g(\tilde{r})\frac{\Gamma(a+b)}{\Gamma(a)\Gamma(b)}\tilde{r}^{a-1}(1-\tilde{r})^{b-1}d\tilde{r}=\int_0^1 g(r)\frac{\Gamma(a+b)}{\Gamma(a)\Gamma(b)}r^{b-1}(1-r)^{a-1}dr. \tag{4.1}$$

Applying Fubini's theorem and substituting $s=r^{-1}$, we have

$$\begin{aligned}&\int_\ell^L f(x)\left\{\frac{\partial}{\partial t}u_t(x)+C_Fx^{\alpha_F}+\mathcal{E}(x)-C_Ip_I(x)\right\}dx\\
=&C_F\mathbb{E}_r\left[\int_\ell^L x^{\alpha_F}f(xr)\mathbb{1}_{xr\geq\ell}u_t(x)dx\right]+C_F\mathbb{E}_r\left[\int_\ell^L x^{\alpha_F}f(x(1-r))\mathbb{1}_{x(1-r)\geq\ell}u_t(x)dx\right]\\
=&C_F\mathbb{E}_r\left[\int_\ell^L x^{\alpha_F}f(xr)\mathbb{1}_{r\geq x^{-1}\ell}u_t(x)dx\right]+C_F\mathbb{E}_{\tilde{r}}\left[\int_\ell^L x^{\alpha_F}f(x\tilde{r})\mathbb{1}_{\tilde{r}\geq x^{-1}\ell}u_t(x)dx\right]\\
=&C_F\mathbb{E}_r\left[r^{-\alpha-1}\int_\ell^{rL}y^{\alpha_F}f(y)u_t(y/r)dy\right]+C_F\mathbb{E}_{\tilde{r}}\left[\tilde{r}^{-\alpha-1}\int_\ell^{\tilde{r}L}y^{\alpha_F}f(y)u_t(y/\tilde{r})dy\right]\\
=&C_F\int_0^1\left\{r^{-\alpha-1}\int_\ell^{rL}y^{\alpha_F}f(y)u_t(y/r)dy\right\}\times\left\{\frac{r^{a-1}(1-r)^{b-1}+r^{b-1}(1-r)^{a-1}}{B(a,b)}\right\}dr\\
=&\int_\ell^L C_Fy^{\alpha_F}\left\{\int_{y/L}^1 r^{-(\alpha-1)}\frac{r^{a-1}(1-r)^{b-1}+r^{b-1}(1-r)^{a-1}}{B(a,b)}u_t(y/r)\times r^{-2}dr\right\}f(y)dy\\
=&\int_\ell^L C_Fy^{\alpha_F}\left\{\int_1^{L/y}\frac{1}{B(a,b)}s^{\alpha-a-b+1}\left((s-1)^{b-1}+(s-1)^{a-1}\right)u_t(ys)ds\right\}f(y)dy.\end{aligned} \tag{4.2}$$

As a result, we have established $\mu_t^\infty(dx)$ as a weak solution to the Equation

$$\frac{\partial}{\partial t}u_t(x)=C_Fx^{\alpha_F}\left\{\int_1^{L/x}\frac{s^{\alpha-a-b+1}\left((s-1)^{b-1}+(s-1)^{a-1}\right)}{B(a,b)}u_t(xs)ds-u_t(x)\right\}-\mathcal{E}(x)u_t(x)+C_Ip_I(x), \tag{4.3}$$

with initial condition $u_0(x) = \frac{1}{L-\ell}$ on $[\ell, L]$.

In particular, if we consider uniform fragmentation processes with $a = b = 1$, the fragmentation ratio is uniform, i.e. $r \sim Uni[0,1]$, and $u_t(x)$ solves

$$\frac{\partial}{\partial t}u_t(x) = C_F x^{\alpha_F}\left\{2\int_1^{L/y} s^{\alpha-1}u_t(xs)\,ds - u_t(x)\right\} - \mathcal{E}(x)u_t(x) + C_I p_I(x), \tag{4.4}$$

with initial condition $u_0(x) = \frac{1}{L-\ell}$ on $[\ell, L]$.

**Remark 4.1.** *One can also construct Equation* (4.4) *by considering $u_t(x)$ as the mass of fragments with length $x$ at time $t$. The integral term on the R.H.S. of* (4.4) *corresponds to the increase in mass at length $x$ due to fragmentation of larger fragments. The $-u_t(x)$ term corresponds to loss of mass at length $x$ due to fragmentation. The last two terms correspond to fragments exiting the system at size-dependent rate $\mathcal{E}(x)$, and new fragments immigrating into the system with a size-dependent rate $C_I p_I(x)$.*

#### 4.2 Stationary solution to FRIME equations

Since the mean measure of the FRIME process can be expressed as a solution to the FRIME Equation in (3.10), and its density (if it exists) satisfies the density equation (4.3), one can study the stationary distribution of the FRIME process with the stationary solution to (4.3).

Let $S(x)$ be a stationary solution to (4.3), then for $x \in [\ell, L]$,

$$\frac{C_F x^{\alpha_F}}{B(a,b)}\int_1^{L/x} s^{\alpha-a-b+1}\left((s-1)^{b-1} + (s-1)^{a-1}\right)S(xs)ds + C_I p_I(x) = (C_F x^{\alpha_F} + \mathcal{E}(x))S(x). \tag{4.5}$$

Furthermore, at stationarity, the rate of mass increase for fragments with size $L$ due to immigration $(C_I p_I(L))$ must offset the rate of mass loss for fragments with size $L$ due to exit $(\mathcal{E}(L)S(L))$ and fragmentation $(C_F L^{\alpha_F})$. Therefore, $S(x)$ must satisfy the boundary condition $S(L) = \frac{C_I p_I(L)}{C_F L^{\alpha_F} + \mathcal{E}(L)}$.

We first consider the case where $a = b = 1$. If we substitute $z = xs$, Equation (4.5) becomes

$$2C_F\int_x^L z^{\alpha-1}S(z)dz + C_I p_I(x) = (C_F x^{\alpha_F} + \mathcal{E}(x))S(x),$$

and taking derivative with respect to $x$ on both sides, we have

$$-2C_F x^{\alpha-1}S(x) + C_I p_I'(x) = (\alpha C_F x^{\alpha-1} + \mathcal{E}'(x))S(x) + (C_F x^{\alpha_F} + \mathcal{E}(x))S'(x),$$

and therefore

$$S'(x) + \frac{\mathcal{E}'(x) + (\alpha_F + 2)C_F x^{\alpha-1}}{C_F x^{\alpha_F} + \mathcal{E}(x)}S(x) = \frac{C_I p_I'(x)}{C_F x^{\alpha_F} + \mathcal{E}(x)}. \tag{4.6}$$

By the variation of parameters method for first order linear differential equations,

$$S(x) = A_1\exp(-P(x)) + A_2\exp(-P(x))\int_\ell^x \frac{C_I p_I'(z)}{C_F z^{\alpha_F} + \mathcal{E}(z)}\exp(P(z))dz, \tag{4.7}$$

where $A_1, A_2 > 0$ is a constant determined by the boundary condition and the integrating factor $P(x)$ is given by

$$P(x) = \int_\ell^x \frac{\mathcal{E}'(z) + (\alpha_F + 2)C_F z^{\alpha-1}}{C_F z^{\alpha_F} + \mathcal{E}(z)}dz. \tag{4.8}$$

Finally, at stationarity, the net mass of fragments with length between $[L - dx, L]$ immigrating into the system must be equal to the net mass of fragments with length between $[L - dx, L]$ leaving the system through fragmentation, so $S(x)$ must satisfy the boundary condition

$$S(L)C_F L^{\alpha_F} + \mathcal{E}(L) = C_I. \tag{4.9}$$

We now consider the shape of the stationary profile for various scenarios of exit mechanisms.

**No Exit Mechanism:** We consider the case where there is only a killing boundary, i.e. $\mathcal{E}(x) = 0$ for $x \geq \ell$ and all fragments with length smaller than $\ell$ dies instantaneously. In this case,

$$P(x) = \int_\ell^x (\alpha_F + 2) z^{-1} dz = (\alpha_F + 2)\log(x) + C_1 \implies \exp(P(x)) = C_2 x^{(\alpha_F+2)}, \tag{4.10}$$

so

$$S(x) = A_1 x^{-(\alpha_F+2)} + A_2 x^{-(\alpha_F+2)} \int_\ell^x \frac{C_I}{C_F} p_I'(z) z^2 dz, \tag{4.11}$$

where $A_1, A_2$ are some constants determined by the boundary condition.

If we further assume that immigrating fragments are uniformly distributed in $[\ell, L]$, the probability density function for the immigration length satisfies $p_I(z) = \frac{1}{L-\ell}\mathbb{1}_{[\ell,L]}(z)$. As a result, the stationary profile satisfies $S(x) = Ax^{-\alpha-2}$. Finally, solving the boundary condition we have $C_I dx = AL^{-\alpha_F-2} \times C_F L^{\alpha_F} dx$, and so $A = \frac{C_I}{C_F} L^2$.

To conclude, the stationary profile is of the form

$$S(x) = \frac{C_I}{C_F} L^2 x^{-\alpha-2}. \tag{4.12}$$

The same result was established in [2] in full and in [3] for the special case where $\alpha = 1$.

**Constant Exit Rate: (CON)** We may also consider the case where $\mathcal{E}(x) \equiv C_E$, then

$$P(x) = \int_\ell^x \frac{(\alpha_F + 2) C_F z^{\alpha_F - 1}}{C_F z^{\alpha_F} + C_E} dz = \frac{\alpha_F + 2}{\alpha_F} \log(C_F x^{\alpha_F} + C_E) + C, \tag{4.13}$$

and so

$$S(x) = A_1 (C_F x^{\alpha_F} + C_E)^{-\frac{\alpha_F+2}{\alpha_F}} + A_2 (C_F x^{\alpha_F} + C_E)^{-\frac{\alpha_F+2}{\alpha_F}} \int_\ell^x C_I p_I'(z) (C_F z^{\alpha_F} + C_E)^{\frac{2}{\alpha_F}} dz, \tag{4.14}$$

where $C_1, A_1, A_2 > 0$ are some constants.

If we consider uniform immigration, then $p_I'(z) \equiv 0$, so the stationary profile is of the form

$$S(x) = A(C_F x^{\alpha_F} + C_E)^{-\frac{\alpha_F+2}{\alpha_F}}, \tag{4.15}$$

with derivative

$$S'(x) = -\frac{\alpha_F + 2}{\alpha_F} A (C_F x^{\alpha_F} + C_E)^{-\frac{2\alpha_F+2}{\alpha_F}} C_F x^{\alpha_F - 1} < 0. \tag{4.16}$$

The stationary profile is therefore strictly decreasing and exhibitis no peak.

**Power Law with Fixed Boundary (PFB):** If we consider $\alpha_F = 1, L = 1$ and take the exit function as $\mathcal{E}(x) = \max\{x^{\alpha_E} - B_E^{\alpha_E}, 0\}$, then for $x < B_E$,

$$\log S(x) = C_1 - \int_\ell^x \frac{3C_F + \alpha_E z^{\alpha_E - 1}}{z^{\alpha_E} + C_F z - B_E^{\alpha_E}} dz, \tag{4.17}$$

where $C_1$ is some constant. A peak is formed at position $p < B_E$ only if $\frac{d}{dx}\log S(x) < 0$ for $x \in [p, B_E]$.

In Figure 3 in the main manuscript, $C_F = 1, B_E = 0.4$. Therefore,

$$\frac{d}{dx}\log S(x) = \frac{3 + \alpha_E x^{\alpha_E - 1}}{x^{\alpha_E} + x - 0.4^{\alpha_E}}.$$

The expression on the right has a root at $x = \frac{1}{3|\alpha_E|}^{\frac{1}{|\alpha_E|}}$. In particular, the root is smaller than 0.4 if and only if $|\alpha_E| < 1.097$. This aligns with the observation that in the PFB panel in Figure 3 in the main manuscript, the peak of the fragmentation profile is clearly to the left of the exit boundary when $\alpha_E = -0.25$.

**Power Law with No Boundary (PNB):** If we consider $\alpha = 1$, and consider the exit function $\mathcal{E}(x) = L^{-\alpha_E} x^{\alpha_E}$. In that case,

$$\begin{aligned} \log S(x) =& C_1 - \int_\ell^x \frac{3C_F + L^{-\alpha_E}\alpha_E z^{\alpha_E - 1}}{z^{\alpha_E} L^{-\alpha_E} + C_F z} dz, \\ \implies S(x) =& C_2 x^{-\alpha_E} (C_F L^{\alpha_E} x^{1-\alpha_E} + 1)^{\frac{3-\alpha_E}{\alpha_E - 1}}, \end{aligned} \tag{4.18}$$

where $C_1, C_2$ are constants determined by the boundary condition.

This gives the best fit line that is very close to simulated data in Figure 3 in the main manuscript.

**Linear Model:** Here we assume that $\mathcal{E}(z) = C_E - m_E z$ and $m_E \neq C_F$ for all $z \leq B_E := \frac{C_E}{m_E}$.

If we take $\alpha = 1$, then

$$P(x) = \int_\ell^x \frac{3C_F - m_E}{C_E + (C_F - m_E)z} dz = -\frac{3C_F - m_E}{C_F - m_E} \log(C_E + x(C_F - m_E)) + C_1, \tag{4.19}$$

for some constant $C_1$,

$$\begin{aligned} S(x) =& A_1 (C_E + x(C_F - m_E))^{\frac{3C_F - m_E}{C_F - m_E}} \\ &+ A_2 (C_E + x(C_F - m_E))^{\frac{3C_F - m_E}{C_F - m_E}} \int_\ell^x \frac{C_I p_I'(z)}{C_F z^{\alpha_F} + \mathcal{E}(z)} (C_E + z(C_F - m_E))^{-\frac{3C_F - m_E}{C_F - m_E}} dz, \end{aligned} \tag{4.20}$$

where $A > 0$ is a constant.

If we assume uniform immigration, i.e. $p_I(x) = 1$, then,

$$S(x) \propto \left( B_E + x \left( \frac{C_F}{m_E} - 1 \right) \right)^{(m_E - 3C_F)/(C_F - m_E)} \tag{4.21}$$

which increases with $x$ in $[0, B_E]$ only if $m_E > 3C_F$. In particular, when $m_E = 2C_F$, the stationary profile $S(x)$ decreases linearly from 0 to $B_E$, and when $m_E = 3C_F$, the stationary profile $S(x)$ is roughly constant in $0, B_E$. This aligns with the observation that $B_E$ and the mode of the length frequencies coincide whenever $m_E > 3C_F$, see Figure 5.

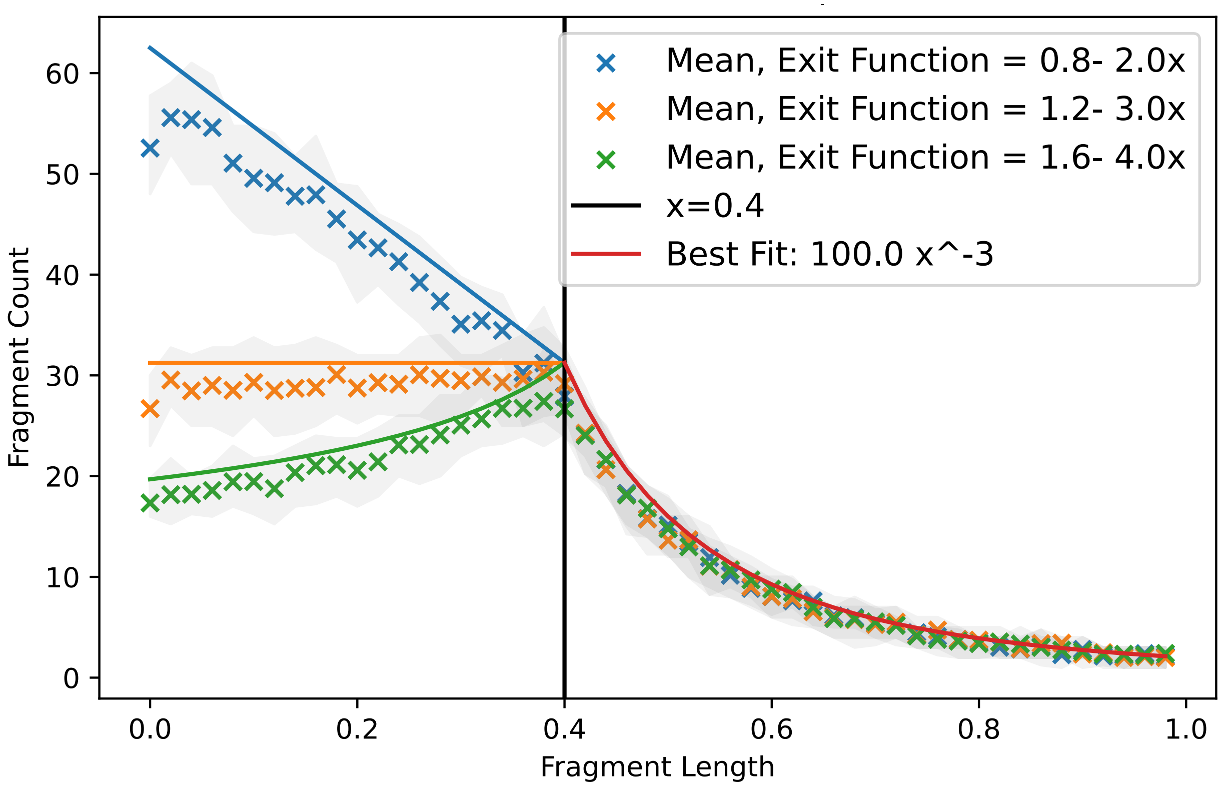


**S9 Fig. FRIME profile with different linear exit mechanism** Stationary distributions of fragment sizes obtained from simulations of the FRIME process with uniform immigration and fragmentation parameter values $C_I = 500, C_F = 1, \alpha_F = 1, a = 1, b = 1$ and simulation threshold $\kappa = 0.05$. 50 points were plotted for each configuration. Each point corresponds to the fragment count per unit length over an interval of length 0.02. Different choices of the immigration function were taken and 50 simulations were run for each choice of immigration function. Shaded regions were plotted using upper and lower quartile fragment count across all simulations. The simulations were run with linear exit mechanism $\mathcal{E}(x) = B_E m_E - m_E x$, where $B_E = 0.4$ and $m_E = 2$ (blue), 3 (orange), 4 (green). A vertical line at $B_E = 0.4$ was plotted in black. A best-fit line of $100x^3$ was plotted in red in the region $x > 0.4$. Note that when $m_E = 4 > 3C_F$, the peak coincides with 0.4 and when $m_E = 2 < 3C_F$, the peak is to the left of 0.4.
